# Synchronization, stochasticity and phase waves in neuronal networks with spatially-structured connectivity

**DOI:** 10.1101/2020.06.04.134940

**Authors:** Anirudh Kulkarni, Jonas Ranft, Vincent Hakim

## Abstract

Oscillations in the beta/low gamma range (10-45 Hz) are recorded in diverse neural structures. They have successfully been modeled as sparsely synchronized oscillations arising from reciprocal interactions between randomly connected excitatory (E) pyramidal cells and local interneurons (I). The synchronization of spatially distant oscillatory spiking E-I modules has been well studied in the rate model framework but less so for modules of spiking neurons. Here, we first show that previously proposed modifications of rate models provide a quantitative description of spiking E-I modules of Exponential Integrate-and-Fire (EIF) neurons. This allows us to analyze the dynamical regimes of sparsely synchronized oscillatory E-I modules connected by long-range excitatory interactions, for two modules, as well as for a chain of such modules. For modules with a large number of neurons (> 10^5^), we obtain results similar to previously obtained ones based on the classic deterministic Wilson-Cowan rate model, with the added bonus that the results quantitatively describe simulations of spiking EIF neurons. However, for modules with a moderate (~ 10^4^) number of neurons, stochastic variations in the spike emission of neurons are important and need to be taken into account. On the one hand, they modify the oscillations in a way that tends to promote synchronization between different modules. On the other hand, independent fluctuations on different modules tend to disrupt synchronization. The correlations between distant oscillatory modules can be described by stochastic equations for the oscillator phases that have been intensely studied in other contexts. On shorter distances, we develop a description that also takes into account amplitude modes and that quantitatively accounts for our simulation data. Stochastic dephasing of neighboring modules produces transient phase gradients and the transient appearance of phase waves. We propose that these stochastically-induced phase waves provide an explanative framework for the observations of traveling waves in the cortex during beta oscillations.

## Introduction

Rhythms and collective oscillations at different frequencies are ubiquitous in neural structures [1]. Numerous works have been devoted to understanding their origins and characteristics [2] which depend both on the neural area and on the activity of the animal. Gamma band oscillations (30-100 Hz) are for instance recorded in the visual cortex as well as several other structures and have been hypothesized to support various functional roles [3]. Beta oscillations (10-30 Hz) are prominent in the motor cortex during movement planning before movement initiation [4] and have traditionally been assigned a role in movement control while more general roles have also been proposed [5].

Several experimental results point out the need to model and analyze the spatial organization of oscillatory activity [6]. An early study [7], using widefield imaging and voltage-sensitive dyes, reported that stimulus-induced oscillatory activity around 10 Hz and 20 Hz was organized in plane waves and spiral waves in the turtle cortex. This spiral-like organization was also reported for pharmacologically induced 10 Hz oscillations in the rat visual cortex [8]. The underlying mechanisms and specific cells involved in the synchronization of two distant regions have more recently started to be investigated using optogenetic manipulations in mice [9].

A motivating illustrative example for the present theoretical study is the observation that beta oscillatory activity during movement preparation exhibits transient episodes of propagating waves in the motor cortex of monkeys [10–13] and humans [14]. These waves appear to propagate along particular anatomical directions with a typical wavelength of 1 cm and a velocity of about 20 cm o s^-1^ [10]. On the structural side, the characterization of the long-range intracortical connectivity in motor cortex have given rise to several quantification endeavors in both monkeys [15,16] and cats [17] with the strength of excitatory responses to microstimulation decaying on a 2 mm length scale. Can one connect these different observations in some ways and how? What is the proper mechanistic interpretation of these observed waves?

The study of waves in oscillatory media has been largely based on the analysis of oscillator synchronisation which itself has a long history [18]. Classic mathematical methods for studying synchronization of weakly coupled oscillators have been extended to study oscillations and travelling waves in spatially extended media in physics and chemistry [19]. In simple descriptions of neural network dynamics based on rate models introduced by Wilson and Cowan [20], also called neural-mass models, the dynamics of a whole set of neurons is reduced to a small set of differential equations approximately describing the temporal evolution of the firing rate of a “typical ‘‘‘neuron in the set. This allows one to directly apply the techniques developed to analyze synchronization of oscillators to study the synchronization properties of sets of oscillating neurons in the rate-model framework. This approach has been followed in a number of works to study the synchronization of spatially-coupled neural networks in the oscillatory regime [21–25]. Rate models with spatially-structured connectivity have also been extensively studied to analyze pattern formation in a neural network context since Amari’s work [26] (see [27–29] for reviews). A limitation is that rate models are generally difficult to quantitatively relate to models of single neurons.

Networks of model spiking neurons provide a more detailed description of neuronal network dynamics than rate models. Studies of networks with random unstructured connectivities have shown the existence of a “sparsely synchronized” oscillatory (SSO) regime [30] in which a collective oscillation exists at the whole population level while spike emission by single neurons is quite stochastic with no significant periodic component. Experimental recordings suggest that neural rhythms in the beta, gamma and higher frequency ranges operate in this regime [2]. Specifically, rhythms in the beta/low gamma range are thought to arise from sparse synchronization between excitatory (E) and inhibitory (I) neuronal populations with reciprocal interactions [31,32], the so-called Pyramidal-Interneuron Gamma (PING) mechanism [33].

A few studies have considered the impact of structured connectivity on sparsely synchronized oscillations in the spiking-network framework. The influence of delays in synaptic transmission was studied in [34] for the synchronization of two neuronal E-I modules oscillating in the high-gamma frequency range, using a classic rate-model formalism. A variety of dynamical regimes was found and qualitative agreement with the bifurcations observed in simulations of a spiking network was reported when the neurons in each E-I module oscillated in a well-synchronized manner. Visually induced gamma oscillations and their dependence on visual stimulus contrast [35] have been investigated by simulations of a spiking network with a two-layer ring-model architecture. Recently, simulations of 2 or 3 coupled E-I modules have been performed [36] to assess how information flow between different modules is correlated with bursts of transient synchrony at gamma frequency.

Spiking networks model more closely biological reality but are more difficult to analyze theoretically. The mathematical analysis of their oscillatory regime is essentially confined to the neighborhood of the oscillatory threshold in parameter space (parameters being the strengths of synaptic connections). General linear stability analyses of E-I spiking networks with spatially structured connectivity have been performed in the absence of transmission delays [37] or taking one into account [38]. Going beyond linear stability is feasible but already somewhat heavy for a single module with unstructured connectivity [30,39]. An exception is the soluble case of deterministic quadratic integrate-and-fire neurons with a wide distribution of frequencies [40] which has been used to study oscillations of an E-I module [41] as well as synchonization between two weakly-coupled modules [42]. Aside from oscillations, the conditions necessary for the existence of a balanced state [43] in a spatially structured network have been studied [44] and it was found in particular that the spatial spread of excitation should be broader than that of inhibitory inputs. Firing correlations have also been studied in such a network state [45]. However, the spatial organization of sparsely synchronized oscillatory activity in spiking networks with spatially structured connectivity remains to be generally studied.

In the present study, our aim is to analyze the synchronization between different local E-I modules, induced by distance-dependent long-range excitation. We focus on the case where each module oscillates in a sparsely synchronized way [30] to model the *in vivo* situation. We combine spiking network simulations in the SSO regime with a mathematical analysis of an “improved” rate model to develop a quantitative picture of the dynamics in such a system. Relating rate models to spiking networks is a classical endeavor. This can be achieved when the collective dynamics is stationary or slowly varying [46], for instance in the presence of slow synapses [47], by an appropriate choice of the f-I curve in the rate model. More recently, different works have shown that mild modifications allow rate models to overcome this limitation of a slow-varying collective dynamics by introducing a timescale that depends of the firing rate. These “adaptive” rate models successfully produced a quantitative description of an uncoupled neuronal population submitted to time-varying inputs [48], as well as of spike synchronization [49] or oscillations driven by spike-frequency-adaptation (SFA) [50] in a recurrently coupled excitatory neural population. Building on this progress, we first develop a rate model with an adaptive timescale [48, 50] and show that it accurately describes population-level oscillations of an E-I spiking neuron module in the SSO regime. This allows us to make use of the large body of work that has been developed to study synchronization in deterministic rate-model equations [23–25]. We show that in spite of the introduction of the adaptive timescale, the model that we use behaves very similarly [21–24] to classic rate models [20]. For two modules, we find that the synchronization between the two-module oscillations depends on the specific pattern of long-range excitation connectivity. Complex dynamical regimes are produced when long-range excitation is weak and targets only excitatory neurons [21, 22], whereas synchronization of the two oscillations of the two modules is otherwise observed. For a chain of oscillatory E-I modules with long-range excitatory coupling decreasing with distance, we similarly find that the connectivity properties of long-range excitation play an important role. In particular, when long-range excitation only targets excitatory neurons, distant modules oscillate with different phases, namely oscillation phase gradients and phase waves spontaneously appear. The bonus as compared to the classical rate-model results, is that, as we show, the obtained results quantitatively agree with simulations of spiking networks of large size.

Simulations of spiking modules of biologically relevant size, of about 10^4^ — 10^5^ neurons each, display significant stochastic variations in module activities that require to go beyond the use of deterministic rate models. Building on previous work [39], we find that finite-size fluctuations can be quantitatively accounted in the adaptive-timescale rate-model framework by adding to it a stochastic component. Even when different modules would fully synchronize in a deterministic context, stochasticity in the module activities produces differences in the phases of oscillation of neighboring modules. [18,19,25]. Classical [19] and more recent results [25,51,52] show how to derive stochastic equations for the oscillatory phases of the different modules that describe stochastic dephasing. In our case, we find that this usual weak-noise reduction to a phase dynamics is accurate only for modules with a very large number of neurons. We obtain a quantitative description of stochastic effects for modules of moderate size, by computing the noise influence on both the phase and amplitude of the module oscillation at the linear level. Stochastic dephasing creates transient bouts of traveling waves in a chain of modules that would be fully synchronized in a deterministic context. We end by obtaining in a chain of E-I modules with long-range excitatory coupling, the probability of dephasing between close E-I modules and the spectrum of phase velocities for the corresponding bouts of traveling waves. We propose that this phenomenon of phase waves induced by stochasticity is at the root of the observation of waves during oscillatory episodes in cortex.

## Results

### Oscillations in an E-I module: rate-model description vs spiking network simulations

We first consider oscillations of neural activity in a local module comprising excitatory (E) and inhibitory (I) neuronal populations with a spatially unstructured connectivity. Oscillations are observed both in rate-model descriptions [20] and in simulations of spiking networks [31–33]. We choose the neurons in our spiking network simulations to be of the Exponential-Integrate-and-Fire type (EIF) [53] (see Eq. (25) for their mathematical definition), which have been shown to describe well the dynamics of cortical neurons [54].

We wish to use a rate model description that quantitatively describes oscillations of the spiking E-I module in the SSO regime [30]. Mild modifications of classic rate models [20] can give quantitatively accurate accounts of simulations of spiking neurons when spike emission has a strong stochastic component, as it is the case in the SSO regime. This was previously demonstrated both for an assembly of independent spiking neurons receiving identical inputs [48] and for spiking neurons coupled by recurrent excitation [50]. Building on these advances, we use the following adaptive timescale rate-model formulation [48, 50] to represent the activity of a group of EIF neurons:

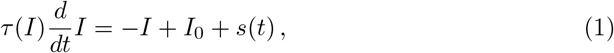

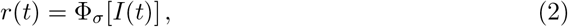

where Φ_σ_(*I*) is the f-I curve of the EIF model (25) for a noisy input current *I_ext_* + *I_syn_* with mean I and total noise strength *σ*. It is shown in Fig. S1A for the parameters used in the present study. This specific choice of f-I curve allows the rate-model to to agree with the rate of a spiking network of spiking EIF neuron in a stationary regime. The term *s*(*t*) represents the time-dependent inputs to the neurons of the population and includes both the external and the network-averaged instantaneous internal inputs. The response time *τ*(*I*) is said to be “adaptive” [48] because it depends on the current *I*. It allows to overcome the limitation of a slowly varying dynamics [46,47] when the population rate remains close to the stationary f-I curve. The chosen function *τ*(*I*) is displayed in Fig. S1B. It is obtained by a fitting procedure to best reproduce the dynamics of a population of uncoupled EIF spiking neurons (see *Methods: Fit of the adaptive timescale in the FAT rate model* for a precise description of our fitting procedure and Fig. S1C for examples of the timescale fit). In agreement with previous works [48,50], Fig. S1D-F shows that the rate model with such a fitted adaptive timescale (FAT model) reproduces quite accurately oscillations of the population activity of a set of identical uncoupled neurons driven by a sinusoidal input current.

The connectivity of our elementary module is depicted in Fig. 1A (inset). It is composed of an excitatory (E) and an inhibitory (I) population with recurrent connections. The FAT rate-model description is obtained by describing the dynamics of each of these populations by Eq. (2),

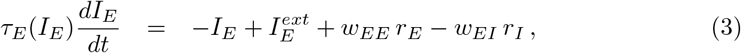

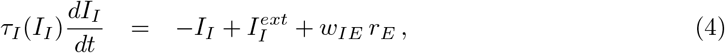

with *r_E_* = Φ_*E*_(*I_E_*) and *r_I_* = Φ_*I*_(*I_I_*). (Here and in the following we have dropped the explicit noise strength index on Φ(*I*) for notational simplicity). In all numerical computations, we have chosen Φ_*E*_(*I*) = Φ_*I*_(*I*) = Φ_*σ*_(*I*), *τ_E_*(*I*) = *τ_I_*(*I*) = *τ*^(*FAT*)^(*I*), with Φ_*σ*_ (*I*) and τ^(*FAT*)^(*I*) displayed in Fig S1. Note that recurrent inhibition has not been included in Eqs. (3,4) both to simplify the analysis and to focus on oscillations in the beta/low gamma range that are mediated by E-I reciprocal interactions. Recurrent inhibitory connections allow for high frequency oscillations in suitable parameters regimes [32,39] and can also be important to prevent a too high firing rate of interneurons, e.g. in the balanced state [43], a role that is assigned here to the external input 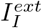.

**Fig 1.**
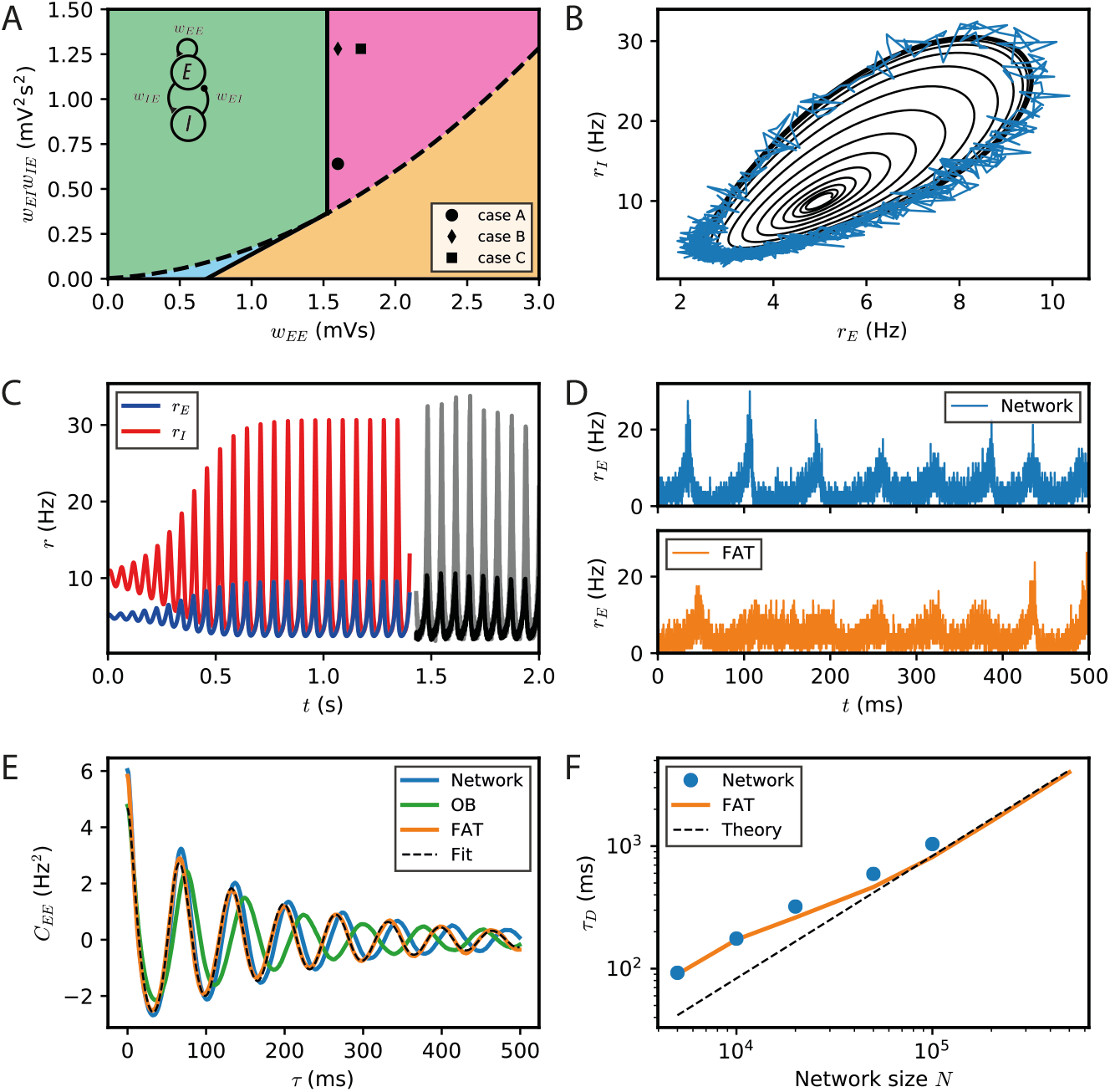
Dynamical regimes of an E-I module. (A) Stability and instability of the module steady discharge as a function of synaptic coupling. Stable regions with complex eigenvalues (green) and real eigenvalues (light blue) and unstable regions with complex eigenvalues (purple) and real eigenvalues (orange) are shown for the stationary state 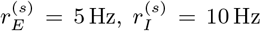. Inset: sketch of the excitatory and inhibitory neuronal populations (the E-I module) and their synaptic interactions. The parameters for the three cases shown are: A – *w_EE_* = 1.6 mVs, *w_IE_w_EI_* = 0.64mV^2^s^2^; B – *w_EE_* = 1.6mVs, *w_IE_w_EI_* = 1.28mV^2^s^2^; C – *w_EE_* = 1.76mVs, *w_IE_w_EI_* = 1.28 mV^2^ s^2^. We use the parameters of case A (solid circle) throughout the remainder of this article; for cases B and C, see Fig. S2. (B,C) Oscillations of the discharge rates *r_E_,r_j_* of the E-I populations in the rate-model description, (B) as *r_I_ vs. r_E_* or (C) as a function of time (blue, excitatory population; red, inhibitory population) for parameters corresponding to case A in panel (A) with *w_IE_* = 2 mVs, *w_EI_* = 0.32 mVs (the corresponding effective constants are *α* = 2.33, *β* = 2.15). Oscillations in a spiking network with a large number of neurons (*N* = 10^6^, resampled with time bin Δ*t* = 0.1ms) are also shown in (B) (blue line) and (C) (black and gray lines). (D) Time traces of the excitatory activity in a spiking network with smaller number of neurons (*N* = 10^4^) and the corresponding stochastic rate model Eqs. (5,6) (sampled with time bin Δ*t* = 0.1ms). (E) Autocorrelation of the excitatory activity for the spiking EIF module (*N* = 10^4^, solid blue) and the adaptive rate model (solid orange). The autocorrelation for the rate model with the adaptive timescale of ref. [48] is also shown (solid green). The fit of the analytical expression Eq. (56) for the autocorrelation to the adaptive rate model (shown in dashed black) allows to obtain the module’s autocorrelation decay time τ_*D*_. (F) The decorrelation time τ_*D*_ shown for spiking modules (solid blue dots), for FAT modules (solid orange line) and as predicted by Eq. (10) (dashed black line).

The stability diagram for the steady non-oscillatory regime of the FAT rate model is easily obtained [20,24] (see *Methods: Oscillatory instability for the E-I module*), see Fig. 1A. As for the classic Wilson-Cowan model [20], it displays three regimes. Steady neural activity is stable when recurrent excitation, measured by the total synaptic strength *w_EE_*, is weak enough. When recurrent excitation grows, two possibilities arise. They depend on the strength of inhibitory feedback on the excitatory population, measured by the product *w_IE_w_EI_*, where *w_IE_* is the total excitatory synaptic strength on inhibitory neurons and *w_EI_* the total inhibitory synaptic strength on excitatory neurons. When inhibitory feedback is weak, the steady state is subject to a non-oscillatory instability: recurrent excitation leads to steady firing at a very high rate, basically limited by the neuron refractory period in our simple model (other mechanisms, not considered here, such as SFA or pair-pulse synaptic depression can moderate this regime). When inhibitory feedback is sufficiently strong, the steady state is destabilized by an oscillatory instability. This instability can lead to finite amplitude oscillations but also to steady high frequency discharge when a steady high-rate fixed point exists (see [55] for a detailed analysis). Oscillations with high discharge and synchronous spiking are also possible outcomes. Here, we limit ourselves to considering oscillations of moderate amplitude that remain sparsely synchronized [30], a dynamical regime which appears most appropriate to describe beta/low gamma oscillations recorded *in vivo.* The parameters corresponding to one such point are indicated in Fig. 1A (solid black circle) and used as reference for the figures of the present paper. The corresponding limit-cycle oscillations are shown in Fig. 1B,C. Two other representative sets of parameter values are also indicated in Fig. 1A (solid losange and square) and the corresponding dynamical traces are shown in Fig. S2.

The rate-model description is compared to simulations of spiking networks (see *Methods: Simulations of spiking networks*) in Fig 1. The rate-model deterministic activity trace accounts well for the network activity when the number *N* of spiking neurons is very large (*N* ~ 10^6^) and stochastic effects at the level of the population are negligible, as shown in Fig 1B,C.

Our E-I module with unstructured connectivity is, however, intended to represent local interactions at a scale comparable to that of a cortical column [56, 57], which is estimated to comprise a smaller number of neurons (*N* ~ 10^4^ — 10^5^). In this case, simulations show that the network population activity has a significant stochastic component, as seen in Fig. 1D. Auto-correlograms of the E population activity display decreasing oscillatory tails reflecting the corresponding dephasing of oscillations (Fig. 1E).

We take into account these stochastic effects in the rate-model description by remembering that, in a sparsely synchronized spiking network [30], the global network activity retains a stochastic component due to the finite numbers *N_E_, N_I_* of excitatory and inhibitory neurons in the network, even for constant external inputs. We follow ref. [39] and assume that, in the SSO regime, the probability of an excitatory (resp. inhibitory) spike between the times *t* and *t* + Δ*t* is given by Φ_*E*_[*I_E_*(*t*)]Δ*t* (resp. Φ_*I*_[*I_I_*(*t*)]Δ*t*), and spikes being independently drawn for each neuron in the network. This leads to replacing the deterministic relations *r_E_* (*t*) = Φ_*E*_ [*I_E_* (*t*)] and *r_I_*(*t*) = Φ_*I*_ [*I_I_*(*t*)] by stochastic versions coming from Poissonian sampling, an approximation that has previously been shown to be quite accurate when neurons are sparsely synchronized [39],

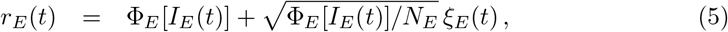

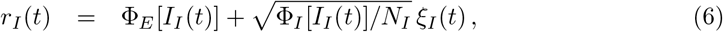

where *ξ_E_*(*t*),*ξ_I_*(*t*) are independent white noises 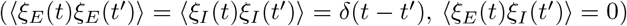 and Ito’s prescription is used [58]. Eq. (5,6) transform the deterministic rate equations into stochastic ones with a noise amplitude that is inversely proportional to the size of the population

Accounting in this way for finite-size fluctuations allows the FAT rate-model description to reproduce quite well the sparsely synchronized oscillations in a moderately-sized spiking network (Fig. 1D). The similarity between the spiking network and the stochastic FAT model can be quantitatively assessed by computing the autocorrelations of the E-I module excitatory population activity, *C_EE_*(*t — t*’) = (*r_E_*(*t*)*r_E_*(*t*’)) — (*r_E_*)^2^. As shown in Fig. 1E, the stochastic FAT model accurately describes the network autocorrelation. It should be noted that the added stochastic terms in the rate equation description have no free parameters, they are entirely determined by the assumption (admittedly approximate) that the instantaneous network spike rate is the product of an underlying Poissonian process.

Stochastic dephasing of oscillations leads to an exponential decrease of the autocorrelation amplitude with a characteristic time *τ_D_*,

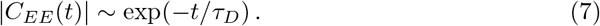

The time *τ_D_* increases with the number of neurons in the E-I module as shown in Fig. 1F.

Well-known results are available to analytically describe dephasing due to weak noise [19,25,51,52], namely in the present case when the E-I module comprises large numbers of neurons and finite-size fluctuations around the deterministic limit cycle are small (see *Methods: Diffusion of the oscillation phase of an E-I module with a finite number of neurons*). For very large numbers of neurons, the activities of the two populations (*r_E_*(*t* + *ϕ*), *r_I_*(*t* + *ϕ*)) are oscillatory with *r_E_*(*t*) and *r_I_*(*t*) periodic functions of time, and the phase *ϕ* being arbitrary but constant. Note that in the present work, we define the phase *ϕ* to be a variable with the same dimension as time and with a period *T* equal to the oscillation period. Another usual convention is to consider the phase as a dimensionless variable with a period of 2*π*. (Therefore *r_E_* is here a periodic function of the phase *ϕ* with period *T*, instead of being a periodic function of a phase with period 2*π*.) The phases in these two different conventions are simply related by a multiplicative factor of 2*π/T*.

Stochastic fluctuations in E-I modules with large number of neurons dominantly produce a random drift of the phase *ϕ* with a diffusive behavior in time,

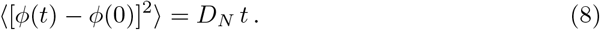

Similarly to the amplitude of the finite-size fluctuations of activity, the diffusion constant *D_N_* vanishes as the numbers *N_E_* of excitatory neurons and *N_I_* of inhibitory neurons in the module grow,

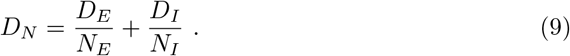

The constants *D_E_* and *D_I_* can be expressed and computed in terms of integrals along the deterministic limit cycle [19] (Eqs. (53,54) in *Methods*). This provides an explicit expression of the decorrelation time as a function of *D_N_* and the period of the limit cycle,

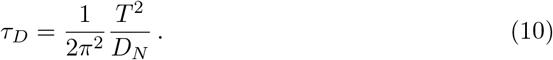

Eq. (10) predicts that the dephasing time increases linearly with the size *N* of the E-I module. Expression (10) is displayed in Fig. 1F. It agrees well with the simulation results for large *N*. Quantitative agreement deteriorates as *N* diminishes and fluctuations become stronger. Fluctuations then strongly affect the shape of limit cycle itself and the description by a pure dephasing becomes less accurate.

### Dynamical regimes of two oscillatory E-I modules coupled by long-range excitation

We start by considering the simplest case of coupling between distant modules, namely two identical E-I modules coupled by long-range excitation. In spite of the addition of the adaptive timescale in the rate model we use, the results we obtain are very similar to results [21,22] obtained for the classic Wilson-Cowan model [20]. It matters for synchronization whether long-range excitation targets only excitatory neurons or both excitatory and inhibitory neurons. We thus distinguish the two cases.

### Long-range excitation targeting excitatory neurons

We analyze the synchronization properties between two E-I modules with oscillatory activity when long-range excitation only targets excitatory neurons, the “*E* → *E*” connectivity case depicted in Fig. 2A.

**Fig 2.**
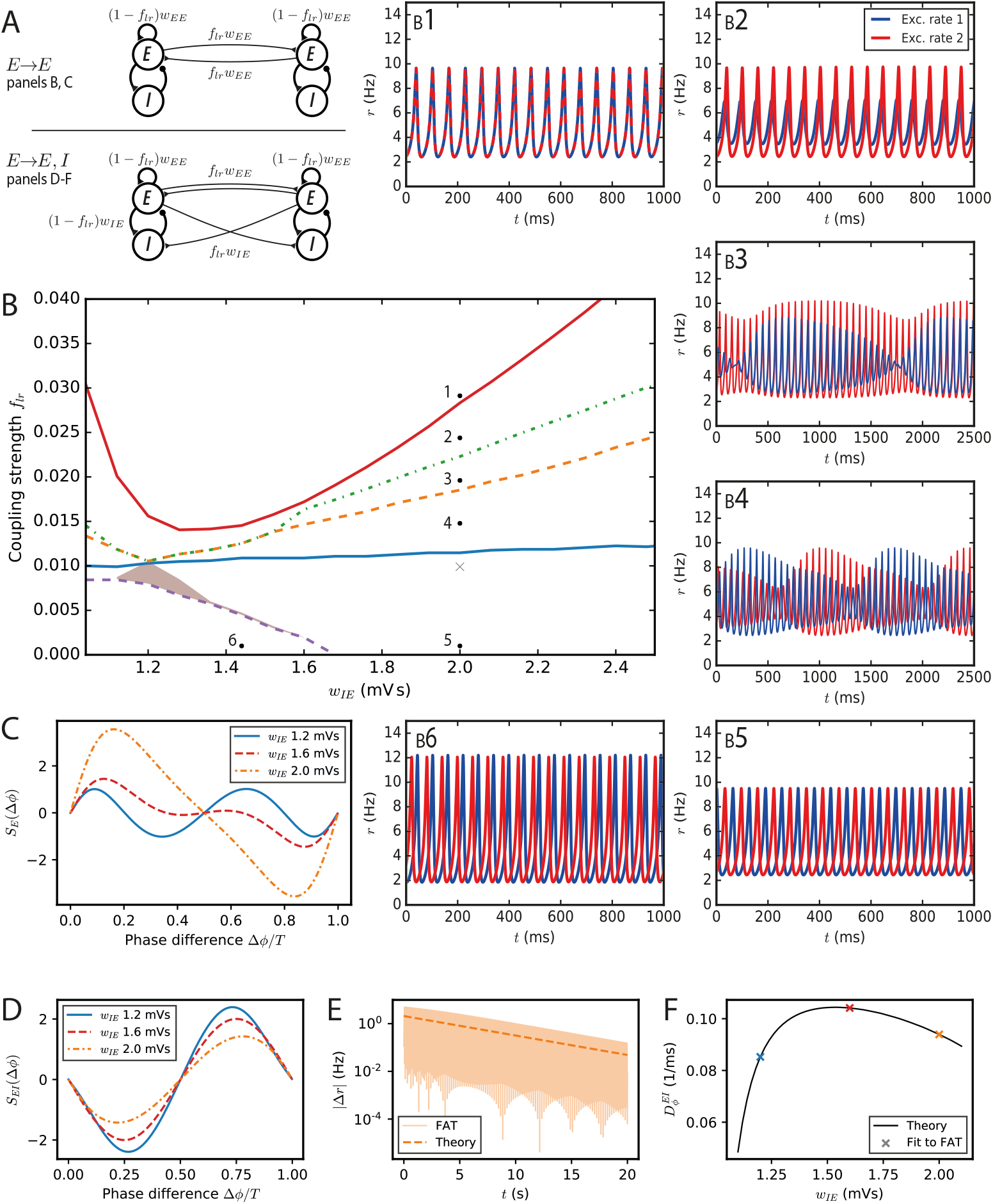
Synchronization and dynamical regimes for two E-I modules coupled by long-range excitation. (A) Sketch of the coupled E-I modules and their synaptic interactions. Top: In the *E* → *E* connectivity case, long-range excitation only targets excitatory neurons in other modules. Bottom: In the *E* → *E, I* connectivity case, long-range excitation targets excitatory and inhibitory neurons in distant modules in equal proportion. (B) Different dynamical behaviors for the coupled modules as a function of the coupling strength *f_lr_* and the synaptic excitatory strength on inhibitory neurons *w_IE_*. The labeled points (solid black circles) mark the parameters of the dynamical regimes shown in (B1-6), the cross (light grey) corresponds to the parameter used in Fig. 3A. (B1) Fully synchronized state for sufficiently strong long-range excitation; *f_lr_* = 0.03. (B2) Finite phase difference: the activity of one module is greater than the other, the amplitudes of their oscillatory activities are constant in time but the phases of their oscillations are different; *f_lr_* = 0.025. (B3) Modulated dominance: one of the two modules is more active than the other but the amplitudes of the oscillatory activities of both modules vary themselves periodically in time; *f_lr_* = 0.02. (B4) Alternating dominance: the activities of the two E-I modules successively dominate; *f_lr_* = 0.015. (B5) Antiphase regime at very low coupling and strong local excitation on inhibitory neurons; *f_lr_* = 0.001. (B6) Finite phase difference regime at very low coupling and weak local excitation on inhibitory neurons; *f_lr_* = 0.001 and *w_IE_* = 1.44 mVs. (C) The synchronization function governing the evolution of the relative of two weakly coupled modules in the *E → E* connectivity case. Modules with weak local excitation of inhibitory neurons *w_IE_* (blue solid line), strong local excitation of inhibitory neurons (orange dashed-dotted line) and *w_IE_* close to the threshold strength of local excitation separating the two regimes (red dashed line). (D) The synchronization function governing the evolution of the relative phase of two weakly coupled modules in the *E → E, I* connectivity case. The fully synchronized state is the only stable state (zero-crossing with negative slope) for the three shown *w_IE_* values. (E) The difference of excitatory activity between the two modules decreases exponentially (*w_IE_* = 2 mVs, *f_lr_* = 0.001). (F) The measured exponential restabilisation in two-modules FAT simulation with *E → E,I* connectivity (crosses, *f_lr_* = 0.001) matches well the prediction of Eq. (15) and Eq. (66) (solid black line).

The dynamics of two coupled E-I modules are described in the rate-model framework (3,4) by

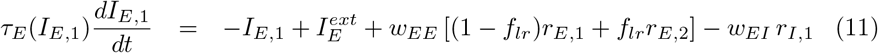

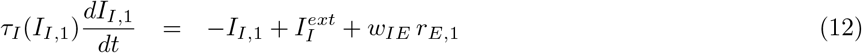

with the same two equations with permuted indices 1 and 2 describing the dynamics of module 2.

We first consider the dynamics of this two-module network for two coupled deterministic FAT rate models, when the firing rates, *r_E,n_* and *r_I,n_* for *n* = 1, 2, are given in terms of the respective currents by the f-I curve (Eq. (2)). Mathematical analysis and simulations show the existence of a number of different dynamical regimes. In order to obtain a precise view of the various cases as a function of the different synaptic couplings, we choose the parameters of the E-I modules so that they remain at a fixed oscillatory location in the parameter diagram of Fig. 1A. This fixes in each module the total strength of recurrent excitatory connections *w_EE_* and the total inhibitory feedback *w_IE_w_EI_*, and leaves as variable parameters the strength of excitation on inhibitory interneurons *w_IE_* (or equivalently the strength of inhibition on excitatory neurons *w_EI_*) and the fraction *f_lr_* of excitatory connections on excitatory neurons that corresponds to long-range excitation. The found dynamical regimes of the two E-I networks are displayed in Fig. 2B as a function of *w_IE_* and *f_lr_* for one of our reference points (case A marked by the black solid circle in Fig. 1A; the analogous diagrams for the two other points are shown in Fig. S3). The individual panels Fig. 2B1–6 show representative traces of the activities of the two excitatory populations in the different regimes. We describe them in turn.

**Fig 6.**
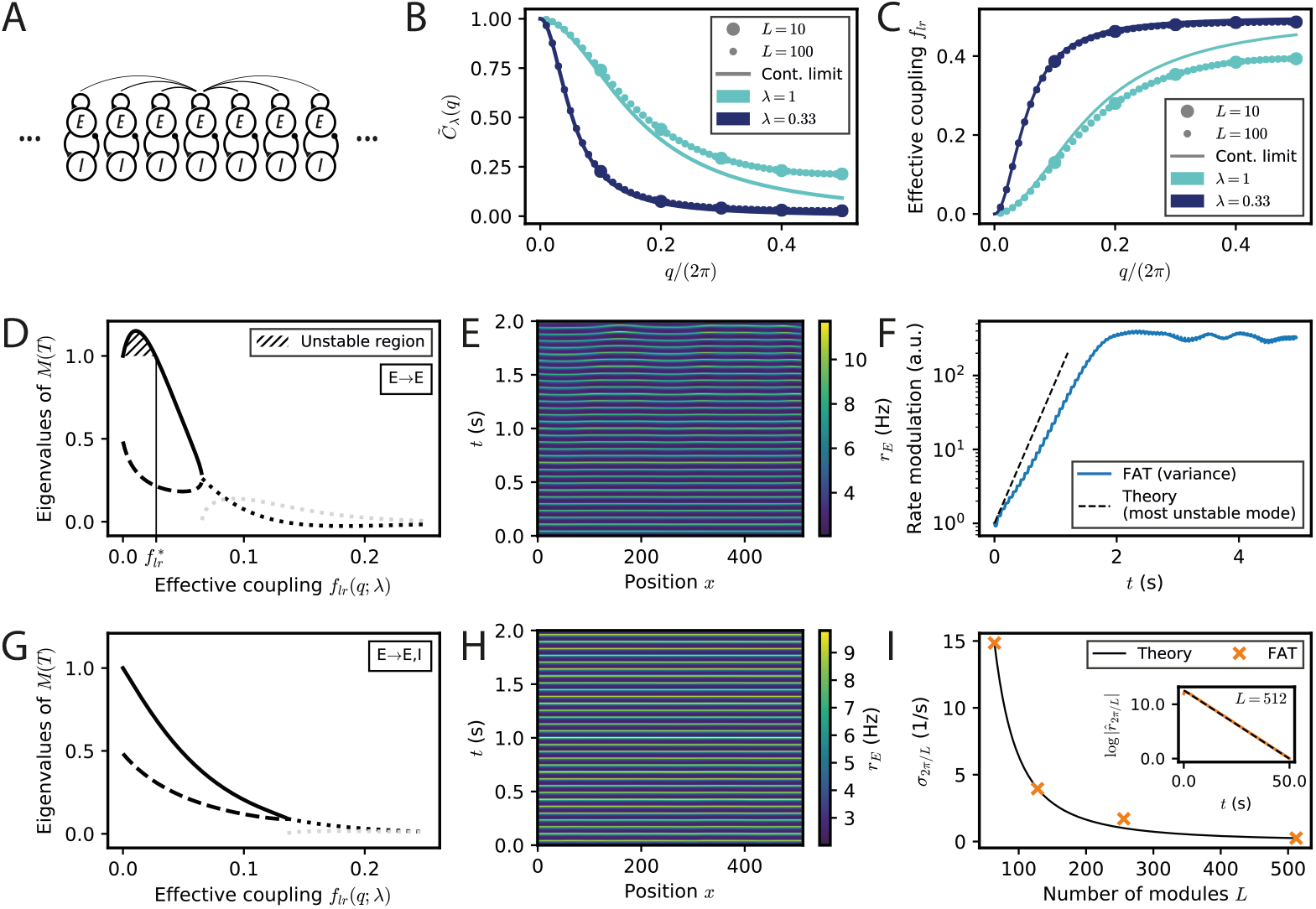
Spontaneous appearance of phase gradients in a chain of E-I modules. (A) Sketch of the chain of E-I modules coupled by long-range excitation. Excitatory neurons in one module target excitatory neurons (*E → E*) or excitatory and inhibitory neurons (*E → E, I*) in the other modules with a distance-dependent excitation profile. (B) Fourier transform of the coupling function 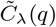 (Eq. (106)) for perturbations of wavenumber *q* when the long-range excitation has an exponential profile (Eq. (20)), for space decay constants λ =1 (turquoise) and λ = 0.33 (dark blue). For λ ≪ 1, the chain of modules can be approximated by a continuous system. (C) Effective long-range coupling 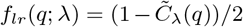 for perturbations of wavenumber *q*. (D) Stability spectrum for perturbations of wavelength *q* as function of the effective coupling *f_lr_* (*q*; λ) for *E → E* connectivity. For small coupling strengths, i.e. long wavelengths, the largest eigenvalue (thick solid line) is larger than 1, and the perturbations are unstable (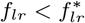, hatched region). While both eigenvalues are real and positive for small coupling strengths, they become complex conjugated for larger couplings (real part, thick dotted black line; imaginary part, thick dotted grey line). (E) Excitatory activity *r_E_* for a chain of deterministic E-I modules over time (*L* = 512, with 0.1 mV noise on initial condition). (F) Growth of the variance of excitatory activity 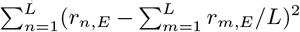 over time, with the predicted rate for the fastest-growing mode shown on top (dashed black line). (G,H) Same as (D,E) but for *E → E, I* connectivity. Note that all modes are stable (G) and no instability develops over time (H). (I) Rate of decay of longest wavelength perturbation *σ*_2*π/L*_ for different chain lengths L (orange crosses) against the theoretical prediction (Eq. (23), black line). Inset: Decay of the spatial modulation of excitatory activity over time (*L* = 512). For all simulations shown, λ = 1/3.

When the E-I modules are not coupled, each of them oscillates with a period *T* and an arbitrary phase with respect to the other module. For very weak long-range excitation, the effect on each E-I module of the excitatory inputs coming from the other one simply leads to a slow change of the phase of its oscillation [19,21,23,24]. The synchronization dynamics itself can be characterized by the evolution of the relative phase Δ*ϕ* between the oscillations of the two modules. For Δ*ϕ* = 0, the two modules oscillate in phase whereas for Δ*ϕ* = T/2 they oscillate in antiphase. The evolution of Δ*ϕ* is found to be governed by the following dynamics [19,21,23,24] (see *Methods: Synchronization function for two weakly coupled E-I modules)*:

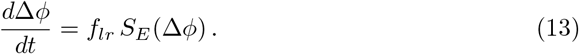

The function *S_E_* (Δ*ϕ*) is shown in Fig. 2C for different synaptic parameter values *w_IE_*. For two identical modules, symmetry implies that the phase differences Δ*ϕ* = 0 and Δ*ϕ* = T/2 are always zeros of *S_E_*. Therefore, in-phase as well as antiphase oscillations are always possible oscillating states. In the present case, the positive slope of the zerocrossing at 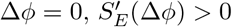 shows that in-phase oscillations are always unstable. When local excitation targets sufficiently strongly interneurons, i.e. for 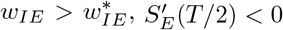 and antiphase oscillations are the only possible stable phase difference. An example of such oscillations is displayed in Fig. 2B5. On the contrary for 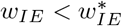, both in-phase and antiphase are unstable. The function S_E_(Δϕ) displays a new zero at a non-trivial Δ*ϕ* with a negative slope (Fig. 2C) and the two modules stably oscillate with this non-trivial phase difference (Fig. 2B6). Such a synchronization regime was also obtained [21] for the classic Wilson-Cowan rate model [20].

For a larger strength of long-range excitation, the interaction between the two modules does not reduce to changing their oscillation phases. Nonetheless, the stability of the inphase oscillations can be computed (see *Methods: Stability analysis of full synchronization for two coupled E-I modules).* The analysis shows that fully synchronous oscillations (Fig. 2B1) are stable when long-range excitation is sufficiently strong. This actually happens for a relatively weak long-range excitation strength, at about *f_lr_* ~ 2 — 7%, depending on the strength *w_IE_* of the local excitation on the inhibitory population in each module (Fig. 2B).

More complex dynamical regimes hold for intermediate strengths of the long-range excitation, below that required for full synchrony and above the excitation strength leading to finite-phase difference phase at very low *f_lr_*. A first decrease of *f_lr_* below that necessary for full synchrony leads to a phase where the two-modules oscillate at the same frequency and different phases. In contrast to the finite-phase difference at very low *f_lr_*, oscillation amplitude in one module strongly dominates the other one, as illustrated in Fig. 2B2. This finite phase difference regime itself looses stability with a further decrease of *f_lr_* and is replaced by a “modulated dominance” regime: beta/low gamma oscillations in one module continue to be of larger amplitude in one module than in the other one but these two unequal amplitudes are themselves modulated at a very low frequency in the 1 Hz range, as shown in Fig. 2B3. This regime stems from a Hopf bifurcation of the finite-difference regime, with the unusual low frequency coming from the small value of *f_lr_* (see *Methods: Stability analysis of full synchronization for two coupled E-I modules*). A further decrease of *f_lr_* transforms the modulated-dominance regime into an “alternating dominance” regime: oscillations in one module dominate the oscillations in the other one, but the dominant module switches periodically, again with a slow frequency in the 1 Hz range, as illustrated in Fig. 2B4. Similarly to modulated dominance, alternating dominance stems from a Hopf bifurcation of the antiphase regime at very low *f_lr_*, when *f_lr_* is increased. A sequence of bifurcations and dynamical states similar to the one reported in Fig. 2 has been described long ago [22] for the Wilson-Cowan rate model upon increase of the excitatory coupling between the two modules. The regimes that we call here “alternating dominance” and “modulated dominance” are referred to as “antisymmetric torus” and “nonsymmetric torus” in ref. [22] and a short discussion of the transition between them is provided.

### Long-range excitation targeting excitatory and inhibitory neurons

We now consider the case in which long-range excitation targets both excitatory and inhibitory neurons with strengths proportional to that of local excitation. In this case, Eq. (12) is replaced by

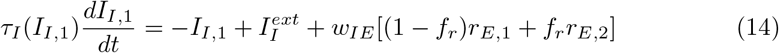

with the same equation with permuted index 1 and 2 to describe the dynamics of the inhibitory population of module 2.

In this “*E → E, I*” connectivity case, sketched in Fig. 2A, long-range excitation is found to always have a synchronizing effect.

For very weak coupling, the synchronization dynamics can be reduced as above to the evolution of the relative phase of the two modules. It is described by Eq. (13) with the function *S_EI_*(Δ*ϕ*) plotted in Fig. 2D. For a small phase difference, *S_EI_*(Δ*ϕ*) behaves as

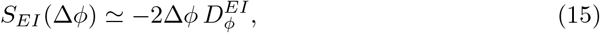

with the constant 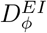 depending on the characteristics of the oscillation cycle (see Eq. (66) in *Methods*). The constant 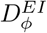 is found to be positive (see Table 1 for its value at the reference point *w_IE_* = 2mVs). Eq. (15) shows that a zero phase lag between the two modules is a dynamically stable configuration, in contrast to our previous results for *E → E* connectivity (compare with Fig. 2C). Eq. (15) predicts that an initial activity difference between two oscillating E-I modules should vanish exponentially. As shown in Fig. 2E, this is indeed observed in simulations with a rate of decrease that agrees well with the one predicted by Eq. (15) (see Fig. 2F).

**Table 1.**
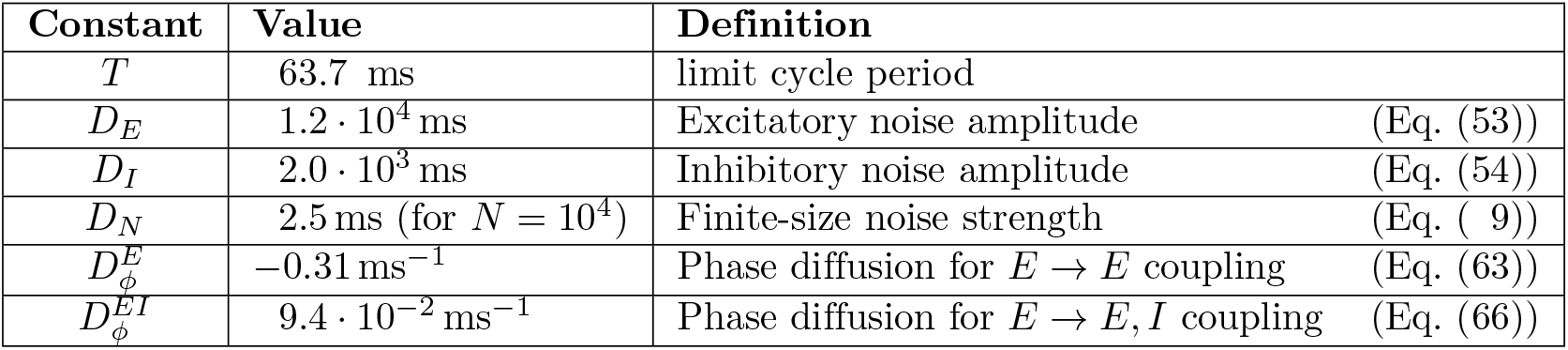
Computed constants. Values of the constants defined in the text for the E-I module limit cycle with the reference parameters (solid circle in Fig. 1A) *w_EE_* = 1.6 mVs, *w_EI_* = 0.32 mVs, *w_IE_* = 2.0 mVs.

| Constant | Value | Definition |
| --- | --- | --- |
| $T$ | 63.7 ms | limit cycle period |
| $D_E$ | $1.2 \cdot 10^4$ ms | Excitatory noise amplitude (Eq. (53)) |
| $D_I$ | $2.0 \cdot 10^3$ ms | Inhibitory noise amplitude (Eq. (54)) |
| $D_N$ | 2.5 ms (for $N = 10^4$) | Finite-size noise strength (Eq. (9)) |
| $D_\phi^E$ | $-0.31 \text{ ms}^{-1}$ | Phase diffusion for $E \rightarrow E$ coupling (Eq. (63)) |
| $D_\phi^{EI}$ | $9.4 \cdot 10^{-2} \text{ ms}^{-1}$ | Phase diffusion for $E \rightarrow E, I$ coupling (Eq. (66)) |

We have checked that this synchronizing effect of long-range excitation persists for stronger long-range excitation strengths. We have found the two-module fully synchronized regime to be linearly stable for all couplings for which we computed its stability matrix (see *Methods: Stability analysis of full synchronization for two coupled E-I modules*). Moreover, numerical simulations with different initial conditions did not show any other stable pattern. Therefore, we conclude that in the context of the present model with instantaneous synapses and no propagation delays, long-range excitation with *E → E, I* connectivity between two E-I modules always tends to fully synchronize their activities.

### Comparison with spiking networks and influence of stochastic activity fluctuations

For a single E-I module, we already found a good match between the oscillatory dynamics in the rate-model framework and simulations of a spiking E-I network, as described in the first section of the results. Finite-size fluctuations were found to play an important role for modules of moderate sizes and were found to be well accounted for by our stochastic rate model. We now pursue this comparison in the two-module case by taking the firing rates *r_E,n_* and *r_I,n_* in Eqs. (11),(12) or (14) expressed in terms of the respective currents by the stochastic relations Eq. (5) and Eq. (6) which account for the finite number of neurons in the networks.

Fig. 3 shows the close correspondence between the rate-model results and networks with a large number of neurons (*N* ~ 10^6^) for *E → E* connectivity. The different dynamical regimes for two E-I modules coupled by long-range *E → E* connectivity are all observed in simulations of networks spiking neurons. Both antiphase (Fig. 3A), alternating dominance (Fig. 3B), modulated dominance (Fig. 3C) and full synchrony (Fig. 3D) are observed for parameters that quantitatively correspond to those predicted by the rate-model analysis. Similarly, in the case of *E → E, I* connectivity, two large E-I modules oscillate in full synchrony as predicted by the rate description (not shown).

**Fig 3.**
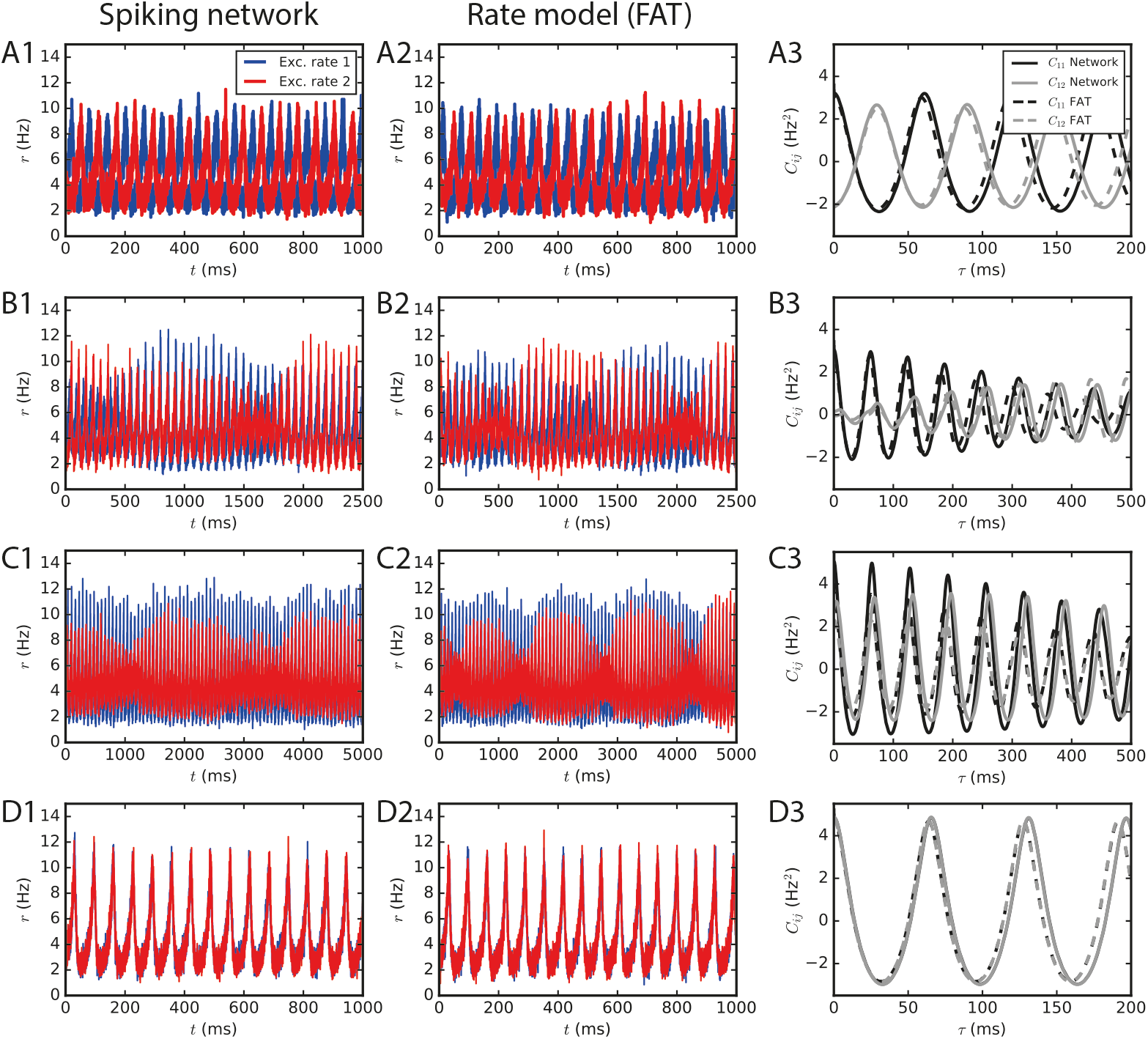
Two large E-I modules coupled by long-range excitation. (1,2) E-I modules with a large number of neurons, *N* = 1.6· 10^6^, and *E → E* connectivity. The different dynamical regimes depicted in Fig. 2 for different coupling strengths are clearly visible both in (1) spiking network simulations and in (2) the corresponding rate-model simulations. The respective auto- and crosscorrelations of the excitatory activities are shown in panels (3). The coupling strengths are as follows: (A) *f_lr_* = 0.01 (light grey cross in Fig. 2B), (B) *f_lr_* = 0.015, (C) *f_lr_* = 0.02, (D) *f_lr_* = 0.05.

For smaller number of neurons, stochastic effects compete with deterministic effects both in simulations of spiking networks and in the rate-model description, when stochasticity arising from finite-size effects is accounted for. Interestingly, both descriptions continue to agree well when fluctuations play a major role in the dephasing of the two E-I modules, as shown in Fig. 4.

For *E → E* connectivity, the correlation between the activities of the two networks clearly displays the signature of the antiphase regime at weak coupling for *N* = 2 · 10^5^ neurons, whereas it is not apparent for *N* = 10^4^ when stochasticity is stronger (Fig. 4A,B). At larger coupling, even small fluctuations lead to modules exchanging their roles and make more complex regimes difficult to identify even for *N* = 2 · 10^5^ neurons. This is illustrated by simulations with a coupling parameter in the modulated dominance regime (*f_lr_* = 0.02, Fig. 4C). The ratio of cross- and auto-correlation functions at zero time lag shows that the transition between different regimes as a function of coupling is blurred by stochasticity as N decreases (Fig. 4E,F).

**Fig 4.**
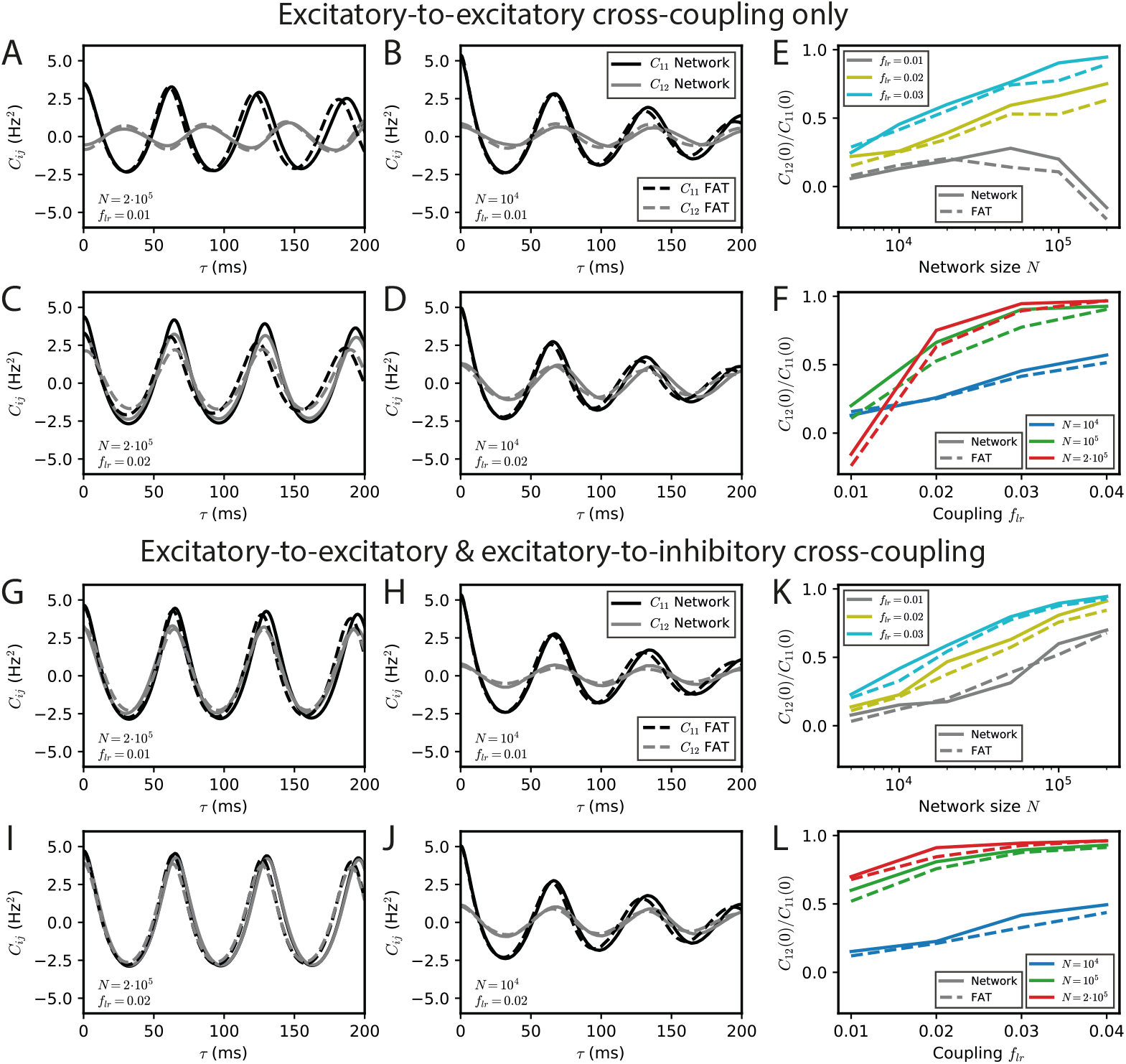
Stochastic effects due to network size for two E-I modules coupled by long-range excitation. Auto- and crosscorrelations *C*_11_ and *C*_12_ of the activities of E-I modules with a finite number of neurons *N* = 10^4^ and 2 · 10^5^ for different coupling strengths *f_lr_* = 0.01 and 0.02 with long-range excitation targeting only excitatory populations (A-D) or targeting both excitatory and inhibitory neurons (G-J) (parameter values are stated in the panels’ lower left corner). Summary plots with the ratio of equal-time crosscorrelations over autocorrelations for varying *N* and *f_lr_* are shown in (E,F) and (K,L), respectively. In the *E → E* case, oscillatory synchrony between the two modules disappears when the coupling strength *f_lr_* diminishes ((C) vs. (D)), but the dynamical regimes of Fig. 2 are blurred by stochastic effects when the network size N decreases ((A,B) vs. (C,D)). In the *E → E,I* case, no other dynamical regimes than oscillatory synchrony are expected, but synchrony increases both with coupling strength flr and, more strongly, with network size *N*.

The corresponding results for E → E, I connectivity are shown in Fig. 4G-L. In this case, two effects can be observed. On the one hand, synchrony decreases at fixed coupling strength when *N* is decreased due to larger noise (Fig. 4G,H). On the other hand, synchronization of the two networks increases when coupling strength increases at fixed *N* (fixed noise) as shown in Fig. 4G,I and Fig. 4H,J. Both effects are quantified in Fig. 4K,L.

One should note that whereas the precise targeting of the long-range excitation plays a crucial role in the instability of in-phase oscillations for large networks (*N* ~ 10^6^), it is much less important for smaller (*N* ~ 10^4^) networks. For these smaller networks, the dominant role of fluctuations in the dynamics of the two individual E-I modules modifies the connectivity synchronization properties. It leads to similar synchronization effects of long-range excitation, whether one module only targets the other E population (*E → E* connectivity) or targets both E and I populations in the other module (*E → E, I* connectivity) (compare Fig. 4B to Fig. 4H).

### Analysis of the competition between synchronization and noise

We thought it interesting to try and analyze more precisely the competition between synchronization and stochastic fluctuations, given that it arises for E-I modules of biologically realistic sizes. We chose to precisely study the case of *E → E, I* connectivity when stochastic fluctuations simply compete with full synchronization promoted by long-range connectivity. We developed two approaches which both assume that fluctuations are of an amplitude moderate enough that they can be treated as perturbations of the deterministic limit cycle.

A classical approach is available for weakly coupled modules and small noise. It was previously used to study noisy coupled oscillators [19] as well as neural systems [25]. In the present case, the outcome of this approach is that finite-size fluctuations in the rate-model description add a stochastic component to the previous Eq. (13) for the phase difference between the two modules (see Eqs. (78) and (84) in *Methods*). (It also provides an estimate to the small shift of oscillation frequency due to small noise [51,52] which we do not consider here). This leads to a very explicit quantification of the difference of oscillation phases between the two modules induced by fluctuations,

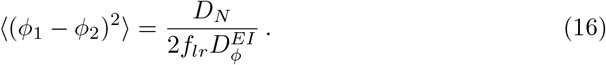

In this expression, the constant *D_N_* is simply the amplitude of stochastic fluctuations in a single module (Eq. (9)), whereas *f_lr_D_EI_* quantifies the synchronization strength due to long-range connectivity (Eqs. (66) and (15)). As intuitively expected, the mean square phase difference between the two oscillators increases with the noise strength and decreases when the synchronization strength increases.

Eq. (16) provides a simple estimate of the competition between stochastic dephasing and synchronization. In order to compare it to numerical simulations, one can compute the respective phase of oscillations of two oscillatory E-I modules. Alternatively, Eq. (16) can be transformed into a prediction of the average spike-rate difference between the two modules, taking into account that the firing rate is a periodic function of the oscillatory phase (Eq. (82) in *Methods).* The latter is chosen to compare with results of numerical simulations of both FAT rate-model and spiking network implementations of two coupled E-I modules in Fig. 5. The mean square difference of excitatory rates between the two modules are quite comparable in the noisy FAT (Fig. 5A) and spiking networks (Fig. 5B), again showing the quantitative accuracy of the FAT rate-model description. However, Fig. 5 shows that the predicted rate difference fluctuations obtained from Eq. (16) (“phase approximation” in Fig. 5) are accurate only for very small coupling between the two modules.

**Fig 5.**
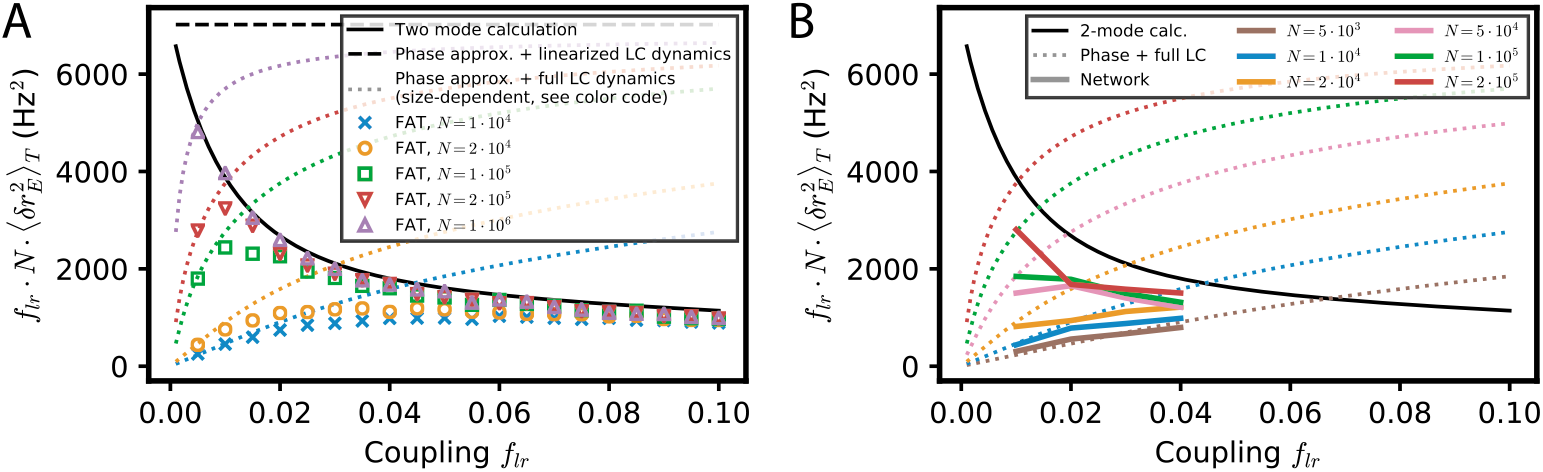
Competition between synchronization and noise for two E-I modules with *E → E, I* connectivity: theory vs. simulations. The variance of excitatory activity between the two modules 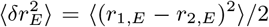 is shown as a function of coupling strength f_lr_. (A) Simulations of the stochastic rate model (FAT, duration 100 s, averaged over 3 repetitions) for different noise strengths (color symbols), as specified in the insert in term of the number of neurons N in the individual E-I module. These simulation results are plotted against theoretical predictions with different approximations. For very small coupling strengths, the phase approximation with the full limit cycle dynamics (Eqs. (16, 82), dotted lines) gives a good account of the observed synchronization, while for larger couplings both phase and amplitude modulations have to be considered (two-mode calculation, Eq. (97), solid black line). Note that for very small couplings, the two-mode calculation coincides with the phase approximation with linearized limit cycle dynamics (Eqs. (16,80), dashed black line), which does not take into account the periodicity of the firing rate as a function of the phase and which predicts an unphysical diverging activity difference for vanishing coupling. (B) Same as in (A) with the results for the network simulations of Fig. 4K,L (colored solid lines corresponding to different E-I module sizes, as sprecified in the inset), shown as 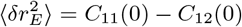.

The underlying assumption in Eq. (16) is that the phase difference plays a dominant role in the difference of activities between the two E-I modules because the phase difference mode has a much smaller restoring strength than the other (amplitude) mode between the two modules. The respective eigenvalues of the two modes are shown in Fig. 6G as a function of the fraction of long-range connection *f_lr_*. Indeed, for weak coupling (*f_lr_* ≪ 1), one eigenvalue 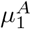 is close to 1, so that the restoring strength 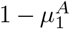 of the corresponding (phase) mode is much smaller than 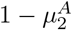, the restoring strength of the other mode. However, an explicit computation shows that 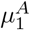 quickly decreases as *f_lr_* increases (see Eq. (76) in *Methods*). Therefore, the two modes soon play comparable roles as *f_lr_* increases.

We developed another approach, valid for small noise but for arbitrary coupling *f_lr_*, in order to check this origin of the inaccuracy of Eq. (16) as well as to obtain a quantitatively precise description of the competition between synchronization and stochasticity. The approach is simply based on accounting for the effect of stochastic perturbations at a linear level around the fully synchronized state where all modules are in the exact same states (see *Methods: Competition between synchronization and stochasticity for two coupled E-I modules of finite size. Weak noise at arbitrary coupling).* As shown in Fig. 5, this two-mode computation produces precise estimates of the mean square excitatory rate difference of the two modules, 〈(*r*_1,*E*_ — *r*_2,*E*_)^2^〉, even for modules with only 10^4^ neurons when stochasticity is quite high. This second approach is thus quite successful in quantitatively capturing the observed competition between synchronization and stochasticity.

### Phase gradients in a chain of oscillatory E-I modules coupled by long-range excitation

Having analyzed the synchronization dynamics and effects of finite-size fluctuations for two E-I modules, we now turn to a fuller description of spatially-structured connectivity. We consider a chain of E-I modules, with each module’s long-range excitation targeting other modules in a distance-dependent way. Interestingly, the effects that we uncovered in the two-module setting, such as desynchronization at weak coupling for *E → E* connectivity and competition of synchronization with stochastic finite-size fluctuations, reappear in a more elaborate form. Moreover, the computational techniques that we developed for the two-module case turn out to be directly applicable to the chain of modules.

### Spontaneous appearance of phase gradients due to long-range excitation specifically targeting excitatory neurons

The setting of our analysis is sketched in Fig. 6A. E-I modules with locally unstructured connectivity are coupled by long-range excitation. For *E → E* connectivity, the dynamics of *L* coupled E-I modules are described in the rate-model framework (3,4) by

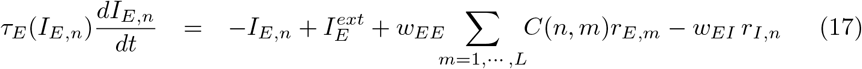

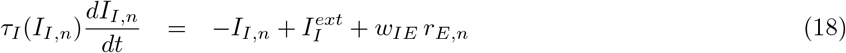

with *C*(*n, m*) = *C*(|*n – m*|).

For *E → E, I* connectivity, Eq. (18) is replaced by

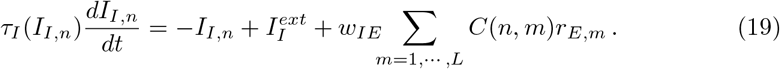

The connection strength decreases monotonically with the distance *l* between modules [17], as described by the function *C*(*l*). Our analytical expressions are valid for a general functional form. For the numerical simulations, we have chosen, for illustration, an exponential decrease which appears compatible with experimental data in the primate motor cortex [16]. We specifically take

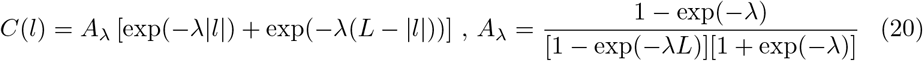

where *A*_λ_ implements the normalization 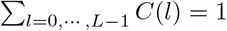. The second exponential corresponds to choosing periodic boundary conditions for the chain. While this is not realistic for the cortex, it minimizes boundary effects in the simulations and should not significantly modify the results in the regime when L is large compared to 1/λ (i.e., when exp(—λ*L*) ≪ 1).

We first analyze the dynamics in the framework of deterministic rate equations, before considering the effects of fluctuations and comparing with spiking networks in the next section.

A perfectly synchronized oscillating chain, with zero phase difference between the oscillations of different modules, is always a possible network state at the level of the rate-model description. It needs however to be tested whether full synchronization is resistant to perturbations or whether some perturbations are amplified by the dynamics and full synchronization is unstable. Interestingly, the linear stability analysis of a periodic lattice such as the considered 1D chain, exactly reduces to that of a two E-I module network with an effective coupling *f_lr_*. More precisely, any small perturbation of the fully synchronized state is the linear sum of periodic perturbations with different wavenumbers q which evolve independently of each other. Perturbations of wavenumber q evolve in the same way as perturbations in the two E-I module network considered in the previous section, provided the coupling in the two-module network is chosen as

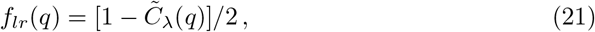

where 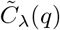 is the Fourier transform of the long-range excitation profile *C*(*l*) (Eq. (106) in *Methods*). 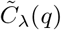 and the corresponding “effective” coupling in the two-module network, *f_lr_*(*q*), are displayed in Fig. 6B,C for an exponentially decreasing long-range excitation profile *C*(*l*) ~ exp(–λ|*l*|). It is seen in Fig. 6C that small wavenumbers *q* correspond to weak effective coupling *f_lr_*(*q*), with *f_lr_*(*q*) → 0 when *q* → 0. While deriving the exact relation (21) requires mathematical analysis (see *Methods: Stability analysis of full synchronization for a chain of oscillatory E-I modules*), it can be intuitively understood. For perturbations of small wavenumbers (*q* → 0), only distant parts of the chain are in distinct oscillating states. The relative dynamics of these distant parts is thus effectively only weakly coupled by excitation since long-range excitation decreases with distance. The relation (21) allows one to directly transcribe for the E-I module chain the results previously obtained for the two E-I module case. As before, we compare two connectivities for long-range excitation.

We first consider the case of *E → E* connectivity, when long-range excitation only targets excitatory neurons. The effective coupling *f_lr_*(*q*) decreases monotonically with the wavenumber *q* and vanishes as *q* → 0. The stability diagram displayed in Fig. 2B then shows that full in-phase synchronization is unstable for chains sufficiently long to accommodate modulations of small enough wavenumber *q*. The associated stability eigenvalues are shown in Fig. 6D. In-phase synchronization is unstable when *f_lr_* becomes smaller than a critical 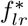 that depends on the single E-I module synaptic parameters. For the the chain of modules, this implies that spatially-periodic perturbations grow when their wavenumber *q* lies below a critical *q*\*, defined by

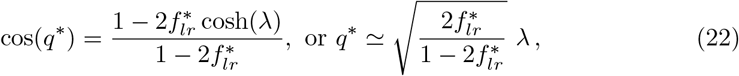

where λ controls the exponential decrease with distance of the long-range excitation. The second equality in Eq. (22) is valid for small λ; the used approximate relation between *f_lr_* and q is shown in Fig. 6C, it can be seen that it is already very accurate for λ = 1/3.

Eq. (22) predicts that for a network of sufficient spatial extent *L* > 2π/*q*^1^, spontaneous phase gradients should appear along the chain. This is confirmed by direct numerical simulations of the rate-model equations, shown in Fig. 6E. The exponential growth of the modulation of excitatory activity along the chain is shown in Fig. 6F, starting from a weakly perturbed fully synchronized state. This directly confirms the linear instability of full synchronization. One further notable feature in Fig. 6E is that the phase gradients develop on a spatial scale that is quite long (i.e. ~ 30 modules) compared to the spatial scale of the long-range excitation profile *C*(*l*) (~ 1 in Fig. 6D). This stems from the fact that in-phase synchronization is unstable only for small *f_lr_* (*q*), that is, for small *q* or long wavelengths compared to the characteristic module coupling length. For our reference parameters and *w_IE_* = 2mVs, Eq. (22) gives for the critical wavelength *q*^1^ ≃ 0.24λ. Similarly, one obtains *q_m_* ≃ 0.15λ for the wavenumber *q_m_* of the fastest-growing mode, which corresponds to the largest eigenvalue in Fig. 6D.

For *E → E, I* connectivity, when long-range excitation targets both excitatory and inhibitory neurons, the previous two E-I module synchronization analysis shows that full synchronization is stable up to vanishingly low coupling *f_lr_*. The stability eigenvalues are shown in Fig. 6G. For all wavelengths, their real part is less than 1. This linear analysis thus predicts that any modulation of activity along the chain should disappear exponentially. This is indeed what is observed in direct simulations of a chain of modules, as illustrated in Fig. 6H. Moreover, our stability analysis shows that the longest wavelength modes are the slowest to vanish. Their relaxation rate is obtained (Eq. (107) in *Methods*) as

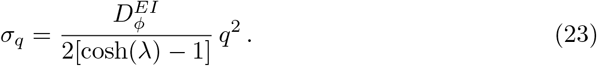

The relaxation rate *σ_q_* vanishes when *q* → 0 and decreases with increasing modulation wavelength. The smallest wavenumber that can be accommodated in a chain of *L* modules is 2*π/L*. Fig. 6I shows that *σ*_2*π/L*_ as given by Eq. (23) precisely describes the approach to full synchronization in our simulations of a E-I module chains of different lengths with long-range *E → E, I* connectivity.

### Phase gradients emerging from stochastic fluctuations in E-I modules of finite size

Spontaneous stochastic fluctuations of activity are present in each E-I module with a finite number of neurons, as already emphasized. In the two E-I module case, these were seen to compete with synchronization and promote phase differences between different E-I modules. Finite-size fluctuations similarly act in a chain of E-I modules. This is shown in Fig. 7, where the results of stochastic rate-model simulations are displayed for both *E → E* and *E → E, I* connectivity. Both the phases of oscillation of different modules (Fig. 7A,D) and the activity of excitatory neurons (7B,E) show that in both connectivity cases differences between neighboring modules are present. The histograms of phase differences between nearest modules are displayed in Fig. 7C,F for *E → E* and for *E → E, I* connectivity, respectively. In both cases, differences in phases decrease when the number of neurons in the modules increases, i.e., when stochastic fluctuations decrease. They also decrease when the range of long-range excitation increases and neighboring modules become more strongly coupled. The similarity between the two connectivity cases shows that stochastic fluctuations play the dominant role in creating phase differences between neighboring modules, for modules of size lower than *N* = 10^5^.

**Fig 7.**
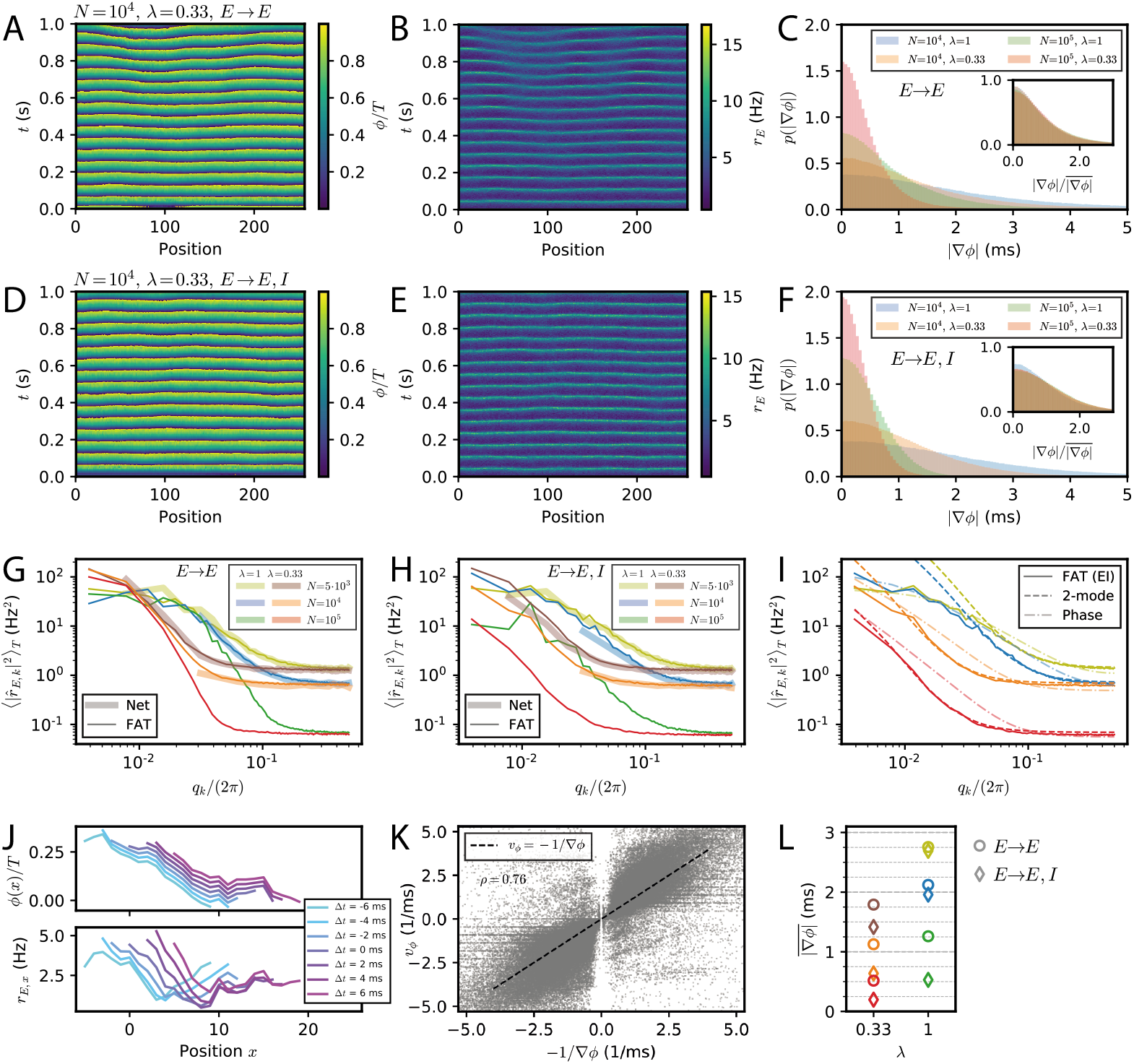
Appearance of phase gradients and stochastic fluctuations in a chain of E-I modules. (A,B) Simulation of the stochastic rate model (FAT) for a chain with *E → E* connectivity, a space constant λ = 0.33 and network size *N* = 10^4^. (A) Instantaneous phase as obtained from the Hilbert transform of bandpass-filtered excitatory activity, where the (unfiltered) rates are shown in (B). (C) Distributions of local phase gradients ∇*ϕ* = *ϕ_x+1_* — *ϕ_x_* for different noise strengths and space constants for *E → E* connectivity. Inset: distributions of gradients normalized to the respective mean value. (D,E) Phase and excitatory activity for the same parameters as in (A,B) but with *E → E,I* connectivity. (F) Same as in (C) but for *E → E, I* connectivity. (G,H) Power spectra of the spatial activity profiles (averaged over 5 s) for different module sizes and space constants λ, for *E → E* (G) and *E → E,I* (H) activity, respectively. The spectra obtained from spiking network simulations (*N* = 5000: *L* = 128; *N* = 10^4^: *L* = 32) are shown in light thick lines, spectra obtained from rate-model simulations are shown in dark thin lines (*L* = 256). (I) Comparison of the spectra obtained from simulations (FAT, solid lines) and theoretical predictions for *E → E, I* connectivity. Similar to the two-module case, the modes with larger effective couplings are well described by the two-mode calculation (dashed lines, Eq. (136)), while for smaller couplings, or vanishing *q*, non-linear effects due to the periodicity of the limit cycle, which are not captured by the linearized two-mode dynamics, become dominant. This limit is better captured by the phase approximation and taking the periodicity of the limit cycle dynamics into account (dash-dotted lines, Eq. (123)). Network size *N* and space constant λ as in (G,H). (J) Example of a spatially extended phase gradient (upper panel) that gives rise to a the apparent propagation from left to right of a (negative) bump of activity in space (lower panel). Phase and excitatory firing rate are shown over a length of 15 modules for different instances in time, see color code legend. (K) Correlation of spatially extended phase gradients 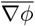 (averaged over 5 modules and retained when the absolute average gradient is larger than the standard deviation of local phase differences) with phase propagation velocities *v_ϕ_* determined from frame-to-frame correlations between spatial phase profiles (see *Methods*). (L) Summary plot of average phase gradients for different network sizes *N* and connection decay space constants λ for *E → E* connectivity (circles) and *E → E,I* connectivity (diamonds).

The stochastic fluctuations along the two chains of E-I modules are further quantified in Fig. 7G,H by computing the power in different spatial modes. These clearly show that for both connectivities, power is highest in the longest wavelengths. This qualitatively agrees with a generalization of the two-module result Eq. (16) for the chain. This result, given in Eq. (127), is obtained by assuming that oscillatory phase differences dominate the fluctuations between modules. While this is true for very long wavelengths, it does not quantitatively agree with the measured spectra for the smaller wavelengths of interest in the present context (Fig. 7I). A generalization of the two-module computation that we developed to take into account both phase and amplitude modes (see *Methods: Characterization of the E-I module chain stochastic activity profile)* is required to quantitatively describe the simulation results, which is also shown in Fig. 7I.

The above results were obtained by simulating chains of E-I modules using the FAT rate-model description. Equivalent simulations for spiking networks are too demanding in computational power for us to perform. We have however checked that for shorter chains of modules (*L* = 128 for *N* = 5000, *L* = 32 for *N* = 10^4^), the stochastic FAT rate model reproduces well the results of spiking network simulations (Fig. 7G,H). This renders us confident that this would be the case for longer chains as well.

### Phase waves and propagation of activity

We have shown that stochastic fluctuations in a chain of E-I modules create phase differences in the oscillations of E-I modules along the chain. In the presence of phase differences, activity in different modules peaks at different times and may produce “phase waves” (also described as traveling waves, e.g. in [2, p1219]), as shown in Fig. 7J. The propagation velocity *v_φ_* of phase waves is kinematically related to the phase gradient and the distance between E-I modules (which is also the linear size of an E-I module),

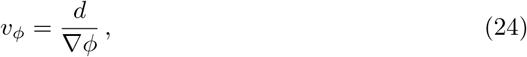

where ∇*ϕ* denotes the phase variation between neighboring modules (recall that the phase has the unit of time with our convention) and *d* is the physical distance between these module centers.

These transient phase waves bear some resemblance to the observation of traveling waves in the motor cortex during episodes of beta oscillations [10–14]. Thus, it appears of interest to better characterize them in the present setting and to compare them to propagation of activity. Phase waves covering a notable spatial range correspond to phase gradients that extend over several modules. Although phase differences between modules are mostly correlated on short length scales, extended phase gradients nevertheless exist. For our purpose, we determined gradients of activity that extend over five modules (see *Methods: Simulations and numerical analysis. Detection of propagation events in an E-I module chain),* and computed the corresponding phase wave velocity. Furthermore, we independently assessed the spatio-temporal propagation of activity for these occurrences by correlating successive frames of simulations (see *Methods: Simulations and numerical analysis. Detection of propagation events in an E-I module chain).* These two independently measured velocities are strongly correlated as shown in Fig. 7K. In our simulations, phase gradients and the associated phase waves thus explain the detected propagation of neural activities.

The probability distributions of spatially extended phase gradients are shown in Fig. S4, for the two different considered connectivities and two space decay constants λ of long-range excitation, as well as for different numbers of neurons in the E-I modules. The corresponding mean values of these gradients are summarized in Fig. 7L. In order to compare these values to neural recordings, values for the size of an E-I module and the range of long-range horizontal connections should be determined. Data are available for different species and for different areas [16,17,59–61]. For instance, for mouse somatosensory barrel cortex, identifying an E-I module with a barrel column would give *d* ≃ 175 *μ*m for the distance between neighboring modules (which is also the linear size of an E-I module) and λ ≃ 0.7 for layer 2/3 horizontal connections since synaptic excitatory charge is measured to decay to 26% two columns away from an excited column [60].

For the motor cortex of adult monkeys, data are available from intra-cortical micro stimulation and recordings with single and multielectrodes [16]. Response amplitudes vary with the protocol and exact location of the electrodes, but a space decay constant of 1.5 mm seems representative. An E-I module size of *d* ≃ 500 *μ*m and *N* ≃ 2 · 10^4^ neurons then gives λ ≃ 0.33 for the long-range connectivity. For this last case, our numerical results give ∇*ϕ* ≃ 0.5ms (*E → E,I*) or 1ms (*E → E*) depending on the precise connectivity (Fig. 7K). With the above estimate of d, Eq. (24) then gives a phase velocity *v_ψ_* ≃ 1 m/s or 0.5 m/s, respectively. In both cases, this is significantly larger than the beta wave velocities recorded in the motor cortex of monkeys. Increasing the local noise on each module increases the dephasing between modules, e.g. ∇*ϕ* ≃ 1.4 ms for *N* =5 · 10^3^ and (*E → E,I*) connectivity (Fig. 7K), and correspondingly lowers *v_φ_*. Thus, in this framework, stochastic dephasing between modules could account for the experimental results only if other sources of “noise” are present beyond the one coming from the finite size of local modules. This would suggest that different E-I modules receive different inputs, shared at the level of individual modules but varying independently from module to module, as further discussed below.

We assessed one other possibility of increasing local noise with respect to the long-range synchronization of neighboring modules. It would consist of diminishing the importance of long-range coupling relative to purely local excitation, without modifying the characteristic length scale of long-range connectivity as such. The associated coupling function then takes the form *C*(*l*) ~ (1 – *α*)*δ*_0*l*_ + αexp(–λ|*l*|), where *α* measures the fraction of truly long-range connectivity. For *α* = 0.5, the resulting average phase gradients are about two times steeper for all *λ* and *N*, see Fig. S5, implying phase waves that are about two times slower than for *α* = 1 (compare Fig. S5E to Fig. 7L).

While noise increase brings the phase wave speed in simulations closer to the range of recorded ones, other features may be responsible for the quantitative differences between the present simplified model and biological recordings, as discussed below.

## Discussion and conclusion

In the present work, we have studied the synchronization of spatially distant E-I modules connected by long-range excitation.

At a technical level, our analysis was greatly facilitated by the formulation of a stochastic rate model with an adaptive timescale that accounts quite precisely for the activity of spiking networks. This extends the finding of previous works [48, 50] to E-I oscillations. The use of a quantitatively precise rate model offers two main advantages. First, it eases numerical simulations that are computationally very demanding for spiking networks with spatially-structured connectivity, since they require both a significant amount of locally connected neurons and a finite number of such modules for spatial extension. Second, the rate-model formulation lends itself much more easily to mathematical analysis [21,24,25,27] than its spiking network counterpart. Therefore, the approach may certainly prove useful in other contexts.

Our application of this methodology provides a description of spatially-structured networks with a local oscillatory dynamics in the SSO regime, based on E-I recurrent interactions. Our first general finding is that, as previously obtained in the Wilson-Cowan rate model framework [21,22,27], the detail of long-range connectivity matters. Based on observations in several neural areas [15–17,61,62], we have confined ourselves to considering long-range excitatory connections. Long-range excitatory connections only targeting distant excitatory neurons result in more complex dynamical properties than long-range excitatory connections that have the same connection specificity as local ones. Experimental investigations of the targeting specificity of long-range excitatory connections appear rather scarce at present (but see [60, 63]). Results such as ours and previous ones [21, 22, 27] will hopefully provide an incentive to gather further data on this question for which experimental tools are now available.

Complex dynamical regimes for two spiking oscillatory modules connected by long-range excitations were previously found [34] for fast oscillations driven by recurrent connections between inhibitory interneurons with synaptic delays, with delays also being present in the excitatory connections. Our results, in agreement with previous rate model calculations [22] extend these previous findings to a different oscillatory regime, for slower beta oscillations, in the absence of recurrent inhibitory connections and significant synaptic delays. It will be interesting to see if these complex regimes are observed in experiments like the one performed in ref. [9] which record and perturb distant neural ensembles oscillating in the beta/low gamma range. They may be also be relevant for oscillations in large ensembles of neurons coupled by long-distance connections, e.g. inter-areal dynamics [64].

We have however observed that these complex oscillatory regimes for two coupled E-I modules are very sensitive to stochastic fluctuations of activity. This has led us to develop a quantitative description of the competition between finite-size fluctuations and synchronization in order to describe the dynamics at the scale of a cortical column. Other studies have proposed more refined and complex ways to take into account finite-size fluctuations (see e.g. [65–67]) than the one we used. But, at least for our purposes, the simple procedure of ref. [39] proved sufficient to account for spiking network simulations.

We have also found that the instability of full synchronization for the two-module network translates in a long-distance instability of synchronization in the extended spatial setting of a chain of modules when long-distance excitation primarily targets excitatory pyramidal cells (*E → E* connectivity). This instability is analogous to those occurring in oscillatory media such as the one created by an oscillatory chemical reaction taking place in a layer of liquid (see [68] for a review). It can similarly be captured by considering the spatio-temporal dynamics of the local oscillation phases as originally proposed in a general coupled-oscillator context [19] and extended in previous works to neural systems [21–25].

This phase instability results in the spontaneous creation of phase differences between neighboring spatial locations, namely oscillatory phase gradients. Phase gradients kinematically lead to the observation of traveling waves of activity (see [2,23] for previous discussions) and seem at the origin of traveling waves associated with oscillatory activity observed in different contexts. The instability wavelength that we determined appears too long however to account for observations in motor cortex [10–14] taking a cortical column typical size of 500 μm and available data on the range of excitatory connections [15–17]. Moreover, this would require that *E → E* connectivity prevails in motor cortex, which is certainly not clear at present.

Stochasticity arising from the finite number of neurons in a cortical column plays a role as important in a spatial setting as for two modules. It competes with the synchronizing effect of connectivity in the *E → E, I* case or in the *E → E* case at length scales smaller than the phase instability critical wavelength. This competition itself induces phase differences between spatially neighboring neural oscillatory ensembles. These produce the stochastic and transient appearance of phase waves, as we have shown (Fig. 7J,K). This has led us to search and obtain a quantitatively precise understanding of stochastic dephasing between different modules. Phase differences are most simply described by stochastic equations for the spatially varying oscillation phase that we derived, and that have been intensely studied in other contexts [69]. We have found however that the stochastic phase description accurately describes oscillatory phase differences only for sufficiently distant spatial locations which appear beyond those relevant for waves within a neural area. On shorter distances, relevant for intra-areal dynamics, modes beyond the phase modes have to be included to obtain a quantitatively accurate description of neural activity and its fluctuations. Dephasing of oscillators in space and the allied phase waves account for the propagating activity events in our simulations (Fig. 7J, K). However, the measured phase wave velocities appear too high to account for the measured velocities [10,12,13] of the transiently appearing traveling waves in the motor cortex during beta episodes. This points to the existence of further effects beyond the ones we have taken into account in the simple model that we have analyzed here. First, there may exist sources of dephasing between modules beyond finite-size noise. It seems plausible that this may arise from inputs from other neural areas which vary from module to module. This seems consistent with the topographic connectivity between the motor cortex and body muscles [15]. Whether these inputs can be treated as white noise or are better approximated by noise with a significant correlation time remains to be seen. Another hypothetic origin of steeper phase gradients suggested by our model would be an overrepresentation of local connections relative to the long-range excitatory connections studied here. This question will have to be addressed quantitatively in light of precise data that will eventually be available. A third non-exclusive possibility is that the slow conduction velocity along non-myelinated axons [59, 62] and the associated propagation delays amplify the phase gradient between different modules. The present analysis provides a solid foundation to incorporate these effects that we plan to examine in forthcoming work together with a detailed comparison with neural recordings.

An interesting further aspect of our results is that the correlation of neural activity at different spatial points is related to the profile of long-range connectivity via the effective coupling. This relation could certainly be tested when the appropriate data becomes available. Interestingly, it could also serve to deduce profiles of long-range connectivity from measurements of neural activity, i.e., measured correlations between spatially distant LFP recordings.

Several other directions will be interesting to pursue to extend the present study. We have here restricted ourselves to consider a one-dimensional chain of E-I modules. Recordings not only display planar waves but also rotating waves and more complex wave patterns. Extension to two dimensions is required for a precise comparison with experimental data [12,13]. Recorded planar waves also display preferred propagation axes [10] that it will be interesting to link to anisotropies in connectivity or possibly to geometrical properties of the motor and premotor cortices. Finally, we have analyzed a very simple E-I module. It will be important to decipher the roles of the different interneuron types (cf. [9]) of the different cortical layers and of their connectivity [60] to better understand cortical dynamics.

## Methods

### Simulations and numerical analysis

#### Simulations of spiking networks

For our spiking network simulations, we used exponential integrate-and-fire (EIF) neurons that exhibit the following membrane potential dynamics:

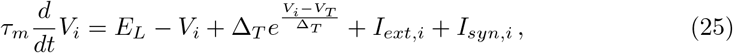

where *τ_m_* = 10 ms is the membrane time constant, *E_L_* = —65 mV is the resting potential set by a leak current, and Δ_*T*_ = 3.5 mV and *V_T_* = —59.9 mV are the sharpness of the spike onset and the spike threshold, respectively. The external current *I_ext_* represents inputs from neurons outside the network and comprises a mean current shared among all neurons of the excitatory or inhibitory population and a noise term specific to each neuron,

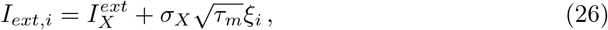

where *ξ_i_* is a Gaussian white noise with zero mean and unit variance, 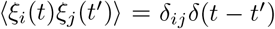, and *X* = *E/I* depending on whether neuron *i* is excitatory or inhibitory. Synaptic currents *I_syn,i_* are given by

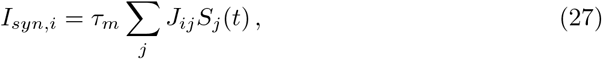

where the *J_ij_* are the synaptic weights between post- and presynaptic neurons *i* and *j*, respectively, and 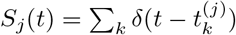 is the spike train of neuron *j* with spike times 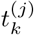.

If not specified otherwise, we considered an all-to-all connectivity at the level of an individual module, such that *J_ij_* ∈ {*J_EE_, J_IE_, J_EI_, J_II_* } depending on the type (excitatory vs. inhibitory) of neurons *i* and *j*. The synaptic weights *J_XY_* relate to population weights *w_XY_* according to *J_XY_* = *w_XY_*/(*N_Yτm_*). Here, *N_E_* and *N_I_* are the numbers of excitatory and inhibitory neurons in a module; for a given network size *N, N_E_* = 0.8N and *N_I_* = 0.2*N*.

We chose the external currents 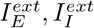 such that at steady state, excitatory and inhibitory neurons fire at an average rate of 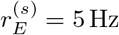 and 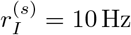, respectively. We furthermore fixed the noise strengths *σ_E_, σ_I_* by imposing a total noise (external plus synaptic) of *σ_tot_* = 10mV, using 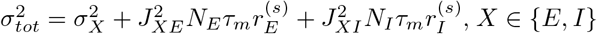. Further parameters were: the numerical threshold potential after which a spike was registered and the membrane potential reset, *V_thresh_* = —30 mV; the reset potential, *V_reset_* = —68 mV; and a refractory period of *τ_ref_* = 1.7 ms. We used a time step dt = 0.01 ms throughout.

For reasons of computational efficiency, we used the following connectivity scheme for the two-module simulations. Because all-to-all connectivity can be implemented without actually processing spikes over all 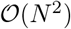 synapses, we maintained all-to-all connectivity with lowered synaptic weights 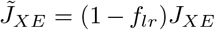 for those synaptic connections within a given module that are supplemented by long-range excitation (*X = E* in the *E → E, X* = *E,I* in the *E → E,I* connectivity case, respectively). The long-range connections were implemented as a finite number *C_lr_* = *f_lr_ N_E_* (rounded to the nearest integer) of excitatory synaptic connections across modules with unchanged synaptic weights *J_XE_*. Only for the very large networks (*N* =1.6 · 10^6^) used in Fig. 3, we also replaced the connections across modules with all-to-all connectivity, where we reduced the synaptic weights accordingly to 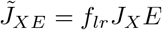.

The long-range excitatory connections between modules in the chain simulations were set up as follows: We first determined the distance-dependent number of excitatory connections as a function of *N_E_* and λ such that the *N_E_* input synapses are distributed according to their relative contribution *p*(Δ*x*) = *e*^-λ|Δ*x*|^/(1 +2/(*e*^λ^ – 1)) across modules, using periodic boundary conditions so that each neuron could in principle receive input from 2 *L*—1 modules. Input connections from different modules were then randomly drawn in corresponding numbers for each neuron receiving long-range excitatory connections.

All network simulations were performed using `brian2`, a convenient Python package for implementing network simulations [70].

Population firing rates were calculated as the total number of spikes within a time bin Δ*t* divided by the size of the population; we typically used Δ*t* = 0.1ms if not explicitly stated otherwise.

#### Computation of correlation functions

The auto- and cross-correlation functions *C*_1,*j*_ (*τ*) with *j* = 1, 2, respectively, as shown in Figures 1–4, were calculated as

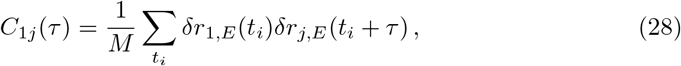

with *M* the number of discrete time points *t_i_* considered, and 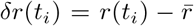 the deviatory part of the firing rate relative to its temporal mean. Note that *M* was limited by the duration of the simulation and we generally discarded an initial transient of 250 ms. We furthermore substracted the expected zero-time Poisson contribution due to stochastic firing, 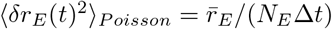, from the autocorrelation at *τ* = 0 to focus on the correlations induced by the network dynamics.

#### Detection of propagation events in an E-I module chain

We first resampled the firing rates obtained by simulations with a new time step Δ*t* = 1 ms by averaging the activity in 1 ms bins to reduce the sampling noise ∝: 1/*dt*. We then determined the Hilbert phase from the band-pass filtered version of the deviatory firing rate 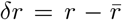, using a fifth-order Butterworth filter with lower and upper corner frequencies of 15 and 40 Hz, respectively. The Hilbert transform was obtained using the Python scipy package.

In a next step, we determined locally averaged phase gradients by averaging phase differences 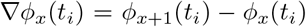 over five neighboring segments, i.e., 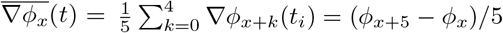 (note that all phase differences are taken modulo a full oscillation period). For the detection of propagation events, we retained gradients where the absolute average phase gradient 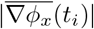 is larger than the standard deviation of the five contributing local phase differences. To determine the propagation of the local phase profile 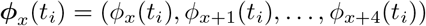, we calculated the Euler distance of laterally shifted versions at later time bins *t_i+j_*,

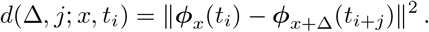

We subsequently inferred the phase propagation speed *v_ϕ_*(*x, t_i_*) from the optimal shift Δ_*opt*_(*j; x,t_i_*) that minimizes the distance between phase profiles after a given time *j*Δ*t* according to:

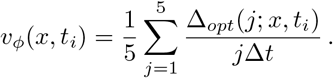

For reasons of computational efficiency, we restricted the search for an optimal shift Δ over shifts of ±15 modules.

### Fit of the adaptive timescale in the FAT rate model

In the adaptive rate model (2), the response time depends on the current *I*. In a previous work [48], Ostojic and Brunel (OB) proposed the analytically-motivated choice,

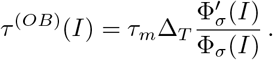

In the present paper, the adaptive response time was chosen following [50], by taking the exponential kernel that best fitted the firing-rate response to an oscillating current at different frequencies. Precisely, we numerically evaluated the analytical expression for the linear firing-rate response 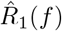 of EIF neurons [71] for frequencies from 1 Hz to 1 kHz in steps of 1 Hz for all values of the input current on the grid of input currents *I* with step size *dI* = 0.1 mV, and fitted the modulus of the firing-rate response by 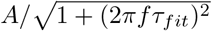, which is the Fourier transformation of an exponential decay in the time domain (see Fig. S1C). We used these values of *τ_fit_* as a look-up table and interpolated in between. The resulting “fitted adaptive timescale” (FAT) curve *τ*^(*FAT*)^ (*I*) is shown in Fig. S1B together with *τ*^(*OB*)^(*I*). We also used look-up tables and interpolation for Φ_*σ*_(*I*) and the first and second derivatives of Φ_*σ*_(*I*) to speed up the numerical calculations. The tabulated functions Φ_*σ*_ (*I*) and *τ*^(*FAT*)^(*I*) used for the present paper are provided as supporting material S1 File.

### Simulations of the FAT rate-model for an E-I module

For a network with synaptic weights *w_EE_, w_IE_* and *w_EI_*, the currents 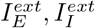 are chosen so as to impose the steady firing rates 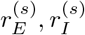 for the excitatory and inhibitory populations. Namely,

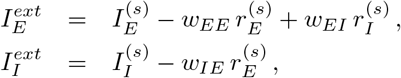

where 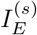 and 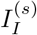 are the currents corresponding to the chosen rates according to 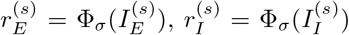. The same external currents are used in the spiking network simulations.

We implemented the stochastic rate-model equations in Python using an Euler scheme with a time step *dt* = 0.01 ms. At each time *t_i_*, we calculated the noisy firing rate *r_X_* (*t_i_*) by drawing a number of emitted spikes *n_X_* (*t_i_*) from a Poisson distribution with mean *N_X_*Φ[*I_X_*(*t_i_*)]*dt*, and setting *r_X_*(*t_i_*) = *n_X_*(*t_i_*)/(*N_X_dt*), *X = E,I*, in order to avoid negative firing rates.

### Oscillatory instability for an E-I module

The E-I module (3,4) either has its E and I populations stably firing at the chosen rates 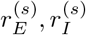 (i.e., the chosen fixed point is stable) or departures from 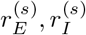 are amplified and the rates cannot be maintained (i.e., the fixed point is unstable). These different cases are shown in parameter space in Fig. 1A. The regions of Fig. 1A are obtained in a standard way [20, 24] by linearizing Eqs. (3,4) around the fixed point currents 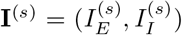 Using a vector notation for the currents, **I** = (*I_E_,I_I_*), the perturbation *δ***I** = **I** — **I**^(s)^ evolves according to

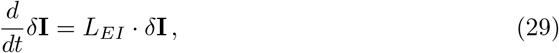

where matrix multiplication is denoted by a dot and *L_EI_* is the 2 × 2 linear stability matrix

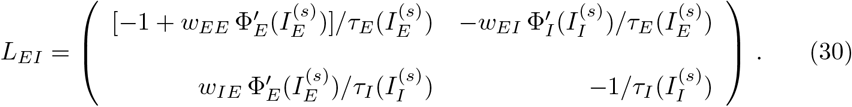

One can note that the derivatives 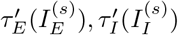 do not appear at the linear level since these timescales multiply time derivatives which are themselves small quantities at the linear level. Thus, the adaptive character of *τ_E_* and *τ_I_* in the FAT model does not modify the computation.

The solutions of the linear system (29) are linear combinations of two exponentials in time, exp(*κt*), with the two arguments *κ* being solutions of

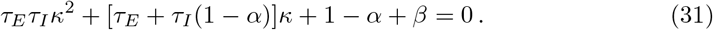

Here, we have defined the two positive constants

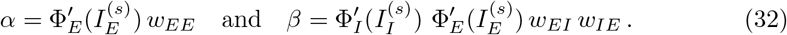

The conditions for the stability of the steady state are that the real parts of the two roots of Eq. (31) are negative, namely that the sum of the two roots is negative and their product is positive, respectively

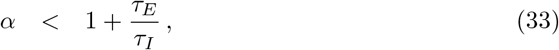

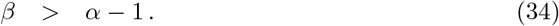

If the latter condition (34) on the product of the two roots is violated, the two solutions are real and of different signs, the instability is purely exponential. When condition (34) holds but not the one on the sum of the two roots (33), the instability is either purely exponential or oscillatory depending on how strongly the second condition is violated. It is oscillatory when *α* — 1 is not too large, namely in the domain

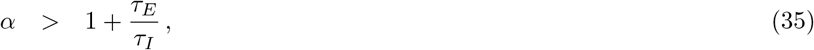

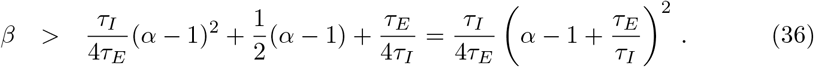

### Phase dynamics of a forced oscillatory E-I module

We consider an E-I module in the oscillatory regime forced by a small time-dependent current, as described by Eqs. (3,4) with

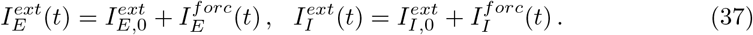

Different choices of the forcing currents, 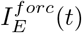 and 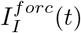, allow us to describe the influence of finite-size stochastic effects as well as the synchronization dynamics of coupled E-I modules. The computation of the drift and entrainment of oscillators is a classical topic [18]. Methods to compute the effect of a weak forcing are well-known [19] and have been applied to neural rate models in numerous works (see [23–25] for reviews). We provide a derivation of these classical formulas for the FAT rate model, for the convenience of the reader.

As above (e.g. Eq. (29)), we adopt a vector notation for the currents. We suppose that the synaptic parameters are such that, with the current 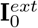 alone, the E-I module follows the limit cycle 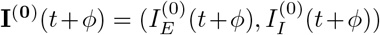 of period *T* (i.e., **I**^(0)^(*t + T*) = **I**^(0)^(*t*)). Here, the phase *ϕ* is an arbitrary constant number reflecting the invariance of the dynamics under time translation when **I**^*ext*^ is constant and there is no time-dependent forcing. When perturbed by a small time-dependent current **I**^*forc*^(*t*), the E-I module currents **I**(*t*) closely follow the unperturbed limit cycle **I**^(0)^(*t* + *ϕ*). However, the phase *ϕ* of the followed limit cycle slowly drifts in time, since it is not submitted to any restoring force.

Note that, for simplicity, in the present work, we define the phase to be a periodic variable between 0 and *T*, instead of normalizing it to 2*π* as it is sometimes done.

In order to determine how **I**^*forc*^(*t*) entrains the phase *ϕ*, we compute perturbatively the correction *δ***I**(*t + ϕ*) to the zeroth-order evolution **I**^(0)^(*t + ϕ*). From Eqs. (3) and (4), we obtain for *δ***I**

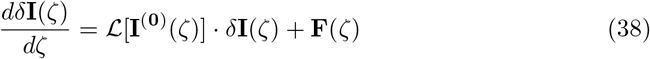

with *ζ* = *t + ϕ* and

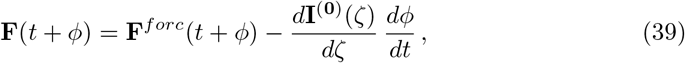

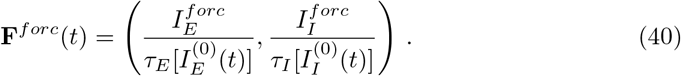

The matrix 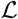 describes the linearized dynamics around the limit cycle,

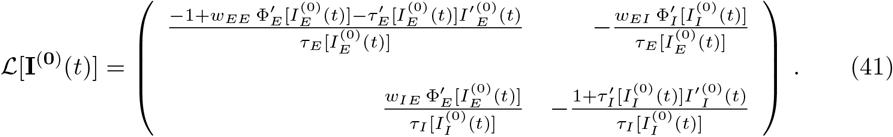

The last term in Eq. (39), proportional to the time derivative of the phase *ϕ*, comes from the slow phase evolution *ϕ* = *ϕ*(*t*) that will be required for preventing the appearance of growing secular terms [72], as explained below.Eq. (38) is a linear equation with coefficients which are periodic functions of time. For this Floquet problem [73], it is helpful to consider the evolution of *δ***I**(*t*) from one period to the next. For this purpose, we introduce the so-called monodromy matrix *M* (*t*) such that

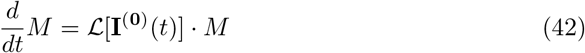

with *M*(0) = *Id*, the identity matrix. The matrix *M*(*t*) describes the linear evolution along the limit cycle and *M*(*T*) the evolution after one full turn around the limit cycle. The vector tangent to the limit cycle 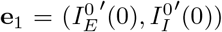 is an eigenvector of *M*(*T*) with eigenvalue *μ*_1_ = 1. The other eigenvector of *M*(*T*), **e**_2_, has an eigenvalue *μ*_2_ with |*μ*_2_| < 1 since the limit cycle is stable. Eq. (38) can be explicitely integrated by writing *δ***I**(*t*) and **F**(*t*) in the basis obtained by translating **e**_1_, **e**_2_ along the limit cycle,

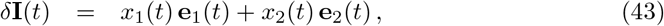

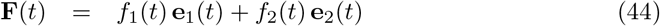

with

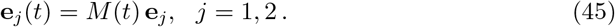

Substitution in Eq. (38) gives

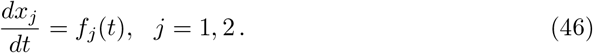

That is after one turn,

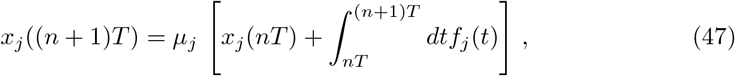

where we have used that *e_j_*(*T*) = *μ_j_e_j_*. When |*μ_j_*| < 1, the above recurrence relation implies that *x_j_* (*nT*) has a magnitude comparable to the integral on the r.h.s. of Eq. (47). In the simple case when *f_j_*(*t*) is periodic of period *T*, the integral is constant and *x_j_*(*nT*) tends toward the finite value 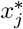,

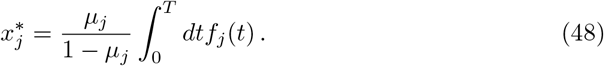

However for *μ*_1_ = 1, Eq. (47) is an arithmetic series and *x*_1_ grows with time, if the integral term in Eq. (47) does not vanish. This would lead to a breakdown of the validity of the perturbation expansion. This growth of the tangent component to the limit cycle reflects a slow phase change of the oscillations that we have anticipated in the last term in Eq. (39). We can now choose the phase evolution so that the integral term in Eq. (47) exactly vanishes and the validity of the perturbative expansion is preserved, requiring that

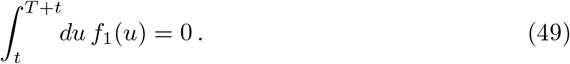

This determines the E-I module phase drift dynamics to be

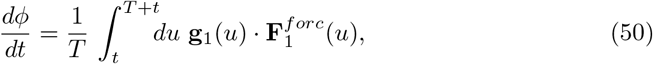

where we have computed the component *f*_1_(*t*) of **F** (Eq. (39)) with the help of the first vector **g**_1_(*t*) = (*g*_1,*E*_(*t*),*g*_1,*I*_(*t*)) of the bi-orthogonal basis (**g**_1_(*t*), **g**_2_(*t*)), such that **g**_*i*_(*t*) · *e_j_* (*t*) = *δ_i,j_*. Eq. (50) can be interpreted in a usual way as the product of the phase response curves of the E-I module multiplied by the perturbations due to forcing [19, 23].

### Diffusion of the oscillation phase in an E-I module with a finite number of neurons

The finite number of spiking neurons in a module gives rise to stochastic effects and produces a diffusive behavior for the oscillation phase of an E-I module in the oscillatory regime. This can be well-captured in the rate-model description by including supplementary stochastic terms, as described by Eqs. (5,6).

These stochastic terms can be interpreted as forcing the deterministic E-I module oscillations. The resultant stochastic phase dynamics can be described by Eq. (50) with **F**^*forc*^(*t*) given by

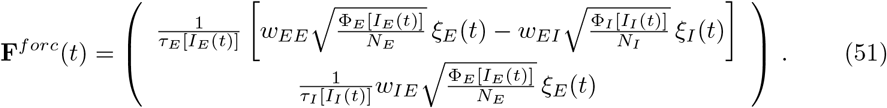

Substitution in Eq. (50) and averaging over the noise gives a diffusive evolution for the phase *ϕ* [19],

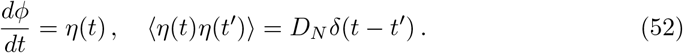

The finite number of neurons in both the excitatory and inhibitory populations contributes to the diffusion constant *D_N_*, as described by Eq. (9). The two constants *D_E_* and *D_I_* in Eq. (9) are expressed as integrals along the limit cycle as

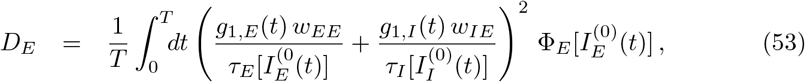

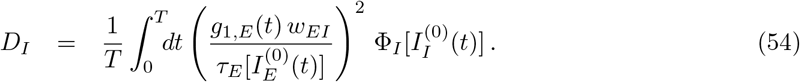

For the synaptic weights *w_EE_* = 1.6mVs, *w_EI_* = 0.32mVs, *w_IE_* = 2.0mVs, Eqs. (53,54) give *D_E_* = 1.2 · 10^4^ ms, *D_I_* = 2.0 · 10^3^ ms (Table 1).

Stochastic effect produces a diffusion of the oscillatory phase, the mean square amplitude of which grows linearly in time, as described by Eq. (8). This gives rise to a loss of oscillation coherence with time and to the decay of the activity correlation functions. Stochastic effects in the two neuronal populations also produce small changes in the oscillation frequency [19,51,52] of magnitudes 1/*N_E_* and 1/*N_I_*, changes that we do not consider here.

The autocorrelation decay time *τ_D_* can be directly related to the phase diffusion constant *D_N_* as follows. We consider, for definiteness, the activity of the excitatory population *r_E_*(*t*). For a very large number of neurons, noise is negligeable and 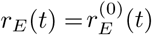 is a periodic function of period *T*. It can written in a Fourier series as

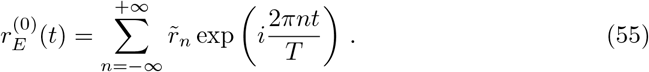

For a finite but large number of neurons, the activity of the excitatory population can approximately be described by taking the dominant effect of phase diffusion into account, namely approximating 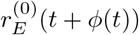. With the help of Eq. (55), it is then easy to compute the auto-correlation of the excitatory activity *C_EE_*(*t*). That is,

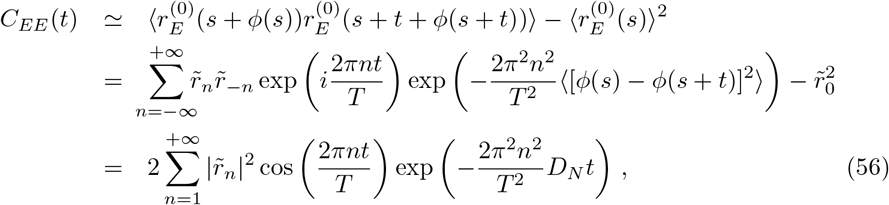

where the angular brackets indicate averages over different realizations of the noise, i.e., the fluctuating values of the phase *ϕ*(*t*). The auto-correlation decay time *τ_D_* is thus obtained as given by Eq. (10). Eqs. (9,10) show that the oscillation coherence time *τ_D_* grows linearly with the number of neurons in the E-I module. For our reference parameters (solid circle in Fig. 1A) where *T* = 64 ms, Eq. (10) gives *τ_D_* ≃ 83 ms for an E-I module with *N* = 10^4^ neurons (*D_N_* = 2.5 ms). For a module with ten times more neurons (*N* = 10^5^), the decay time is ten times longer.

A fit of the activity auto-correlation decay with the above expression allows one to directly measure *τ_D_* in simulations. This procedure is followed in Fig. 1F to compare the theoretical prediction to the simulation results.

### Synchronization function for two weakly coupled oscillatory E-I modules

The phase dynamics of a single forced oscillatory module (Eq. (50)) allows us to derive Eq. (13) governing the evolution of the respective phases of two weakly coupled oscillatory modules [19,21,24,25].

We suppose that the parameters are such that for a vanishing coupling *f_lr_* (Eqs. (11,14)), the two E-I modules follow the limit cycle 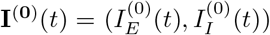 of period *T* with different phases *ϕ*_1_ and *ϕ*_2_,

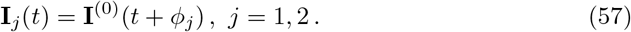

In order to determine how a small coupling *f_lr_* makes the phases *ϕ*_1_ and *ϕ*_2_ evolve, we compute perturbatively in *f_lr_* the corrections *δ***I**_*j*_(*t + ϕ_j_*), *j* = 1, 2, to the zeroth-order evolution (57). We obtain that *δ***I**_1_ follows Eq. (38) with the following forcing current (Eq. (37)) that arises from the coupling to the second E-I module,

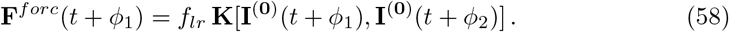

The perturbation comprises both long-range excitation coming from the other module and missing local excitation within the module, which is instead used as long-range excitation onto the other module.

More precisely, for *E → E* connectivity (Eqs. (11,12)), the forcing current is given by

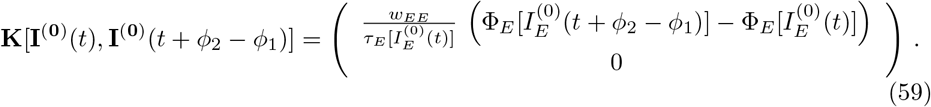

Eq. (50) then determines the phase drift of the first E-I module to be

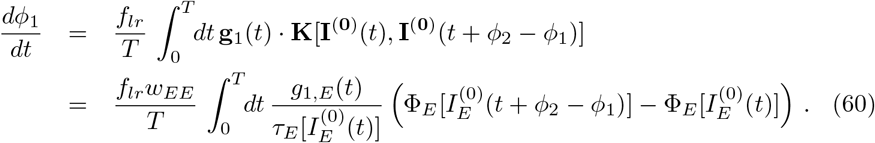

The evolution of *ϕ*_2_ is obtained by permutation of *ϕ*_1_ and *ϕ*_2_ in Eq. (60). Subtraction of these two equations provides the evolution of the relative phase Δ*ϕ* = *ϕ*_1_ – *ϕ*_2_ of the two modules as given by Eq. (13), with the explicit expression of the coupling function *S_E_* being

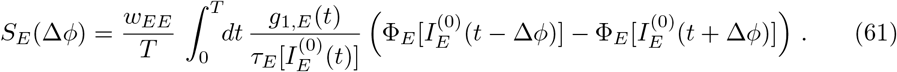

The subscript *E* of *S_E_* denotes that the expression holds in the case when long-range excitation only targets excitatory neurons.

For a small phase difference, one readily obtains to first order in Δ*ϕ*

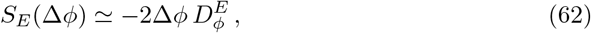

with

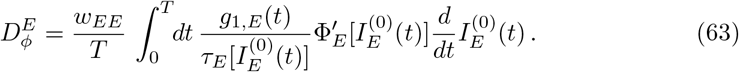

The constant 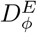 is actually negative (see Table 1 for our reference parameters) so that full oscillatory synchrony is unstable for small long-range excitatory coupling *f_flr_* between the two modules.

For *E → E, I* connectivity (Eqs. (11,14)), the coupling function **K** of Eq. (59) is replaced by

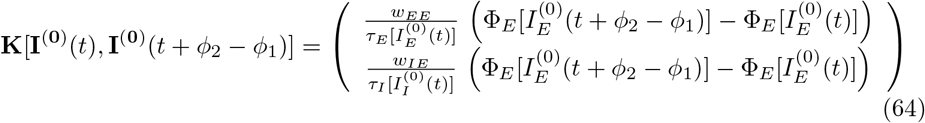

This gives in turn the coupling function for the phase difference Δ*ϕ* between the two modules,

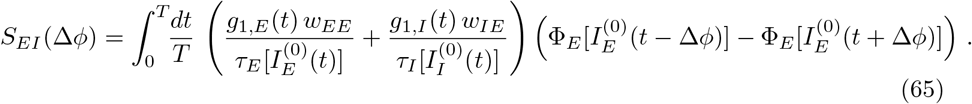

For a small phase difference between the two modules, analogously to Eq. (61), Eq. (65) reduces to Eq. (15), the analog of Eq. (62) for *E → E, I* connectivity, with

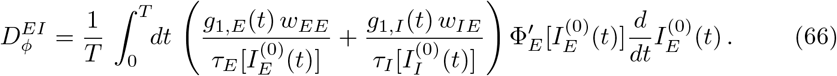

In this case, the constant 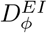 is positive (see Table 1 for our reference parameters) and full oscillatory synchrony is stable for small long-range excitatory coupling between the two modules.

### Stability analysis of full synchronization for two coupled E-I modules

The fully synchronized oscillatory state is always a possible dynamical state for two coupled identical E-I modules. Namely, **I**_1_(*t*) = **I**_2_ (*t*) = **I**^(0)^(*t*) is an exact solution of the coupled Eqs. (17) and (18/19) since the missing local excitation is supplied by the distant modules. The stability of this oscillatory state can be determined irrespective of the strength of coupling between the two modules.

We consider slightly perturbed evolutions for the two modules **I**_*j*_ (*t*) = **I**^(0)^(*t*) + *δ***I**_*j*_ (*t*), *j* = 1, 2. The dynamical behavior of the perturbations *δ***I**_*j*_ (*t*) is obtained by linearizing the dynamics around the fully synchronized oscillatory state,

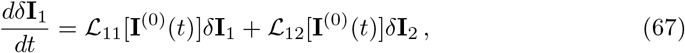

together with the analogous equation with permuted indices 1 and 2.

Since the dynamics is invariant under the interchange of module 1 and 2, the 4-dimensional linear evolution of (*δ***I**_1_(*t*), *δ***I**_2_(*t*)) can be reduced to the study of the 2-dimensional evolution of symmetric (*δ***I**_1_ = *δ***I**_2_) and antisymmetric (*δ***I**_1_ = —*δ***I**_2_) perturbations. The evolution of symmetric perturbations is governed by the 2 × 2 matrix 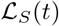 which is identical to the matrix 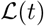 (Eq. (41)) for a single E-I module,

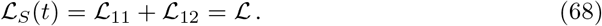

The coupled system of two modules thus inherits its stability with respect to symmetric perturbations from the stability of the limit cycle of a single E-I module. On the contrary, antisymmetric perturbations evolve according to the new 2 × 2 matrix 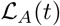,

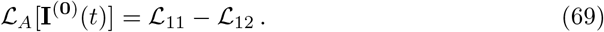

The matrix 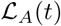 explicitely depends on the coupling *f_lr_*. The stability of the synchronized state is determined by assessing whether antisymmetric perturbations have grown or decayed after one period *T*. As above, this is governed by the eigenvalues of the monodromy matrix *M_A_*(*T*) associated with 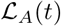,

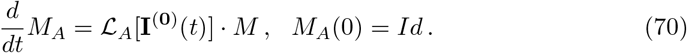

For *E → E* connectivity (Eqs. (11,12)), the matrices 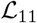 and 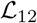 read

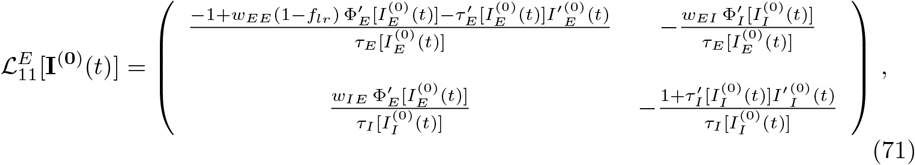

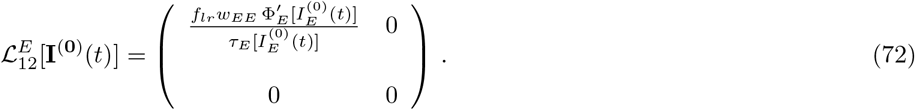

Integration of Eq. (70) for different *f_lr_* and other parameters fixed, shows that the two eigenvalues of 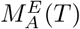 are smaller than 1 for *f_lr_* above a threshold coupling 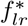, while one eigenvalue is larger than 1 for 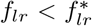. The obtained 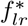 is shown as a solid red line in Fig. 2B.

For *E → E,I* connectivity (Eqs. (11,14)), the calculation is identical but the resulting matrix 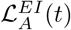 reads

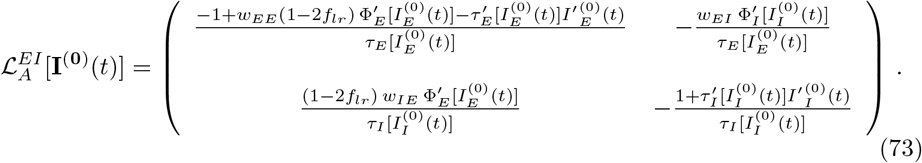

The eigenvalues of the associated monodromy matrix 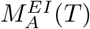 are found to be smaller than 1 irrespective of the value of *f_lr_* (see Fig. 6G), showing that the fully synchronized state is stable in this case.

In the previous section, it was shown that the stability of the synchronized state for small *f_lr_* was dependent on the sign of the constants 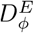 (Eq. (63)) and 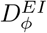 (Eq. (66)), respectively. This can also be seen directly by computing the largest eigenvalue 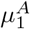 of *M_A_*(*T*) (Eq. (70)) for small *f_lr_* as we shall demonstrate below.

When *f_lr_* = 0, the two modules are not coupled and *M_A_*(*T*) reduces to *M*(*T*), the matrix for a single module (Eq. (42)), with 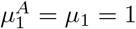. For small 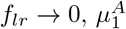 can be computed perturbatively. We write the expansion of 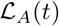 and *M_A_*(*T*) in powers of *f_lr_* to first order as

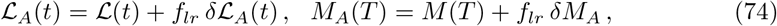

where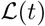 is given by Eq. (41) and the matrix *M* by Eq. (42). The matrix *δM_A_* is obtained from the integration of Eq. (70) to linear order in *f_lr_*,

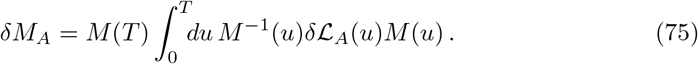

First-order perturbation theory gives the development of the eigenvalue 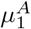 as a function of *δM_A_*,

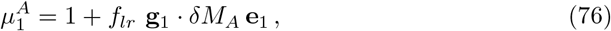

where **g**_1_ the left eigenvector of *M*(*T*) associated to the eigenvalue 1, such that **g**_1_ ° **e**_1_ = 1 (i.e., the first vector of the bi-orthogonal basis (**g**_1_, **g**_2_)). Explicit formulas are obtained after the replacement of *δM_A_* by its expression (75) together with the identity **g**_1_(*u*) = **g**_1_M^-1^ (*u*) and the expressions of 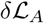 for the two considered connectivities. For *E → E, I* connectivity, Eq. (73) gives 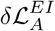, and subsequently the expression for the largest eigenvalue of 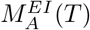,

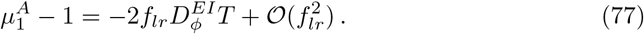

The same formula with 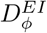 replaced by 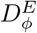 is obtained for the largest eigenvalue of 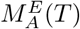 with the help of Eqs. (71) and (72). Since 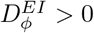 and 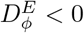, one recovers that *E → E, I* connectivity synchronizes the two modules’ oscillations 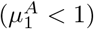 while full synchronization is unstable 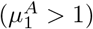 at small coupling for *E → E* connectivity.

### Competition between synchronization and stochasticity for two coupled E-I modules of finite size

For *E → E, I* connectivity, long-range excitation between two E-I modules tends to synchronize their oscillations. In contrast, finite-size stochasticity tends to make their oscillatory phases drift independently. The competition between these two effects can be analytically estimated when noise is weak. We first consider the simplest case when the coupling between the two modules is also weak, before considering the case of an arbitrary coupling.

#### Weakly-coupled modules

The phase dynamics of weakly-coupled modules is simply described by the linear addition of the stochastic diffusion of the two modules’ phases (Eqs. (8,52)) and synchronization due to long-range coupling (Eq. (13)). Thus, one obtains for the phase difference dynamics

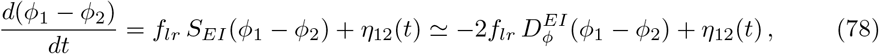

where in the second approximate equality we have replaced *S_EI_*(Δ*ϕ*) by its linear approximation for small phase differences (Eq. (15)). The noise term *η*_12_(*t*) is the difference of the independent finite-size noises for the two modules (52). Being a linear combination of white noises, *η*_12_(*t*) is also a white noise, with an amplitude twice as large as for a single module,

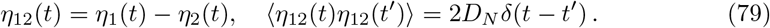

Eqs. (78) and (79) readily give that for a small noise amplitude 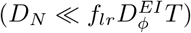, the two-module oscillation phase difference has Gaussian fluctuations with a mean square amplitude given by Eq. (16).

We can use that result to estimate the mean square amplitude of the excitatory firing rate difference 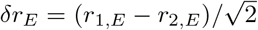 between the two modules. In the case of weak coupling, the firing rates can be approximated as 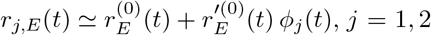, as long as |*ϕ_j_*| ≪ *T*. The firing rate fluctuations are then directly obtained from the phase differences as

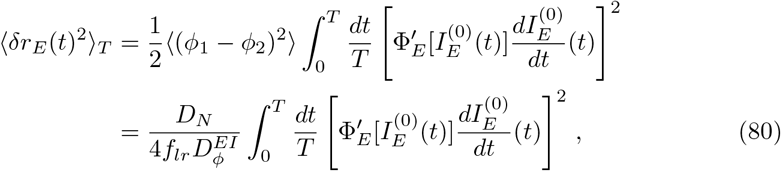

where we used 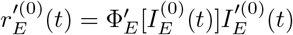 and furthermore averaged over one oscillation period, indicated by the subscript *T*. Note that we neglected here the additional direct, i.e. not phase-mediated, contribution due to Poissonian sampling, which would add an additional term 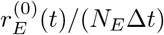 with Δ*t* being the spike sampling interval.

The above expression diverges with vanishing *f_lr_*. This arises from the fact that the phase difference between the two oscillating modules becomes arbitrarily large when the inter-module coupling vanishes. However a large phase difference does not imply large rate differences since for completely uncorrelated oscillating modules the rate fluctuations are given by 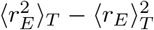. The unphysical divergence of expression (80) for small *f_lr_* comes from the neglect of the periodicity of the rate as function of the oscillation phase. We can obtain an expression for 〈*δr_E_*(*t*)^2^〉_*T*_ that takes this periodicity into account. Using the Fourier decomposition Eq. (55) and 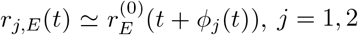, one obtains analogously to the single module autocorrelation

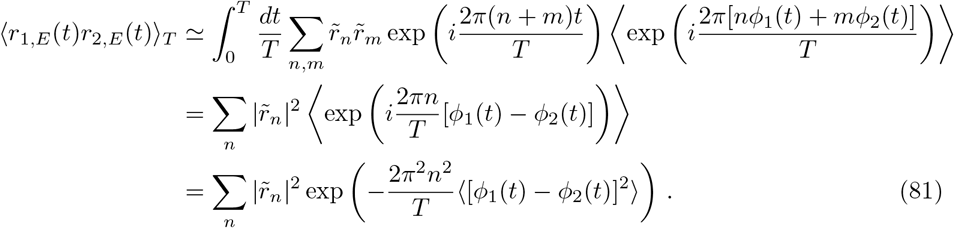

Here, we first used that 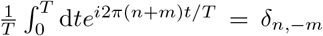 and that the relative phases *ϕ*_1_(*t*), *ϕ*_2_ (*t*) vary slowly with time with respect to the oscillatory dynamics, i.e., can be considered constant over one oscillation period.

Using 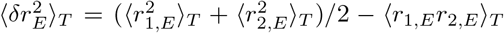, it is now straightforward to obtain

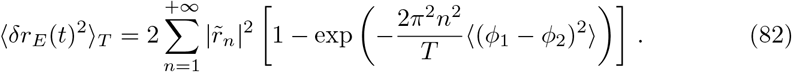

A comparison of this theoretical result with simulations of the stochastic FAT rate model and spiking networks is shown in Fig. 5.

#### Weak noise at arbitrary coupling

Eq. (16) provides a simple estimate of the competition between stochastic dephasing and synchronization for weak coupling between the two modules. A more precise estimate for small noise but arbitrary coupling is obtained by considering general fluctuations around the fully synchronous state that result from the modules’ stochastic activities. Namely, we describe the dynamics around the fully synchronized state, at the linear level, as in Eq. (67), **I**_*j*_(*t*) = **I**^(0)^(*t*) + *δ***I**_*j*_(*t*), *j* = 1, 2, and compute the dynamics of the perturbations *δ***I**_*j*_ (*t*) resulting from the presence of stochasticity. For the first module, this gives

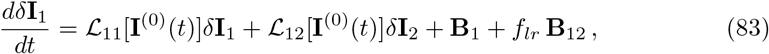

with the stochastic terms

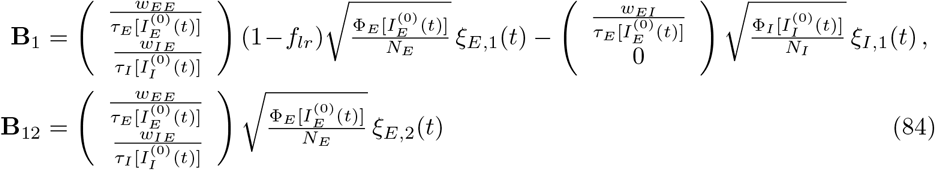

Permuting indices 1 and 2 in Eqs. (83) and (84) gives the dynamics of the linear departure *δ***I**_2_ of the second module from the fully synchronized state.

As previously, the permutation symmetry between module 1 and 2 can be taken advantage of to reduce the 4-dimensional dynamics to a pair of uncoupled two-dimensional dynamics for the symmetric and antisymmetric modes. We focus on the antisymmetric mode *δ***I**_*A*_(*t*) in the following,

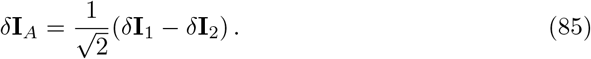

From Eq. (83) and the analogous equation for module 2, the antisymmetric mode obeys

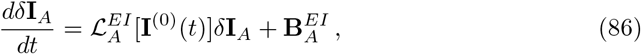

with the matrix 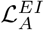 for *E → E, I* connectivity given by Eq. (73). The stochastic term 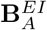 is given by

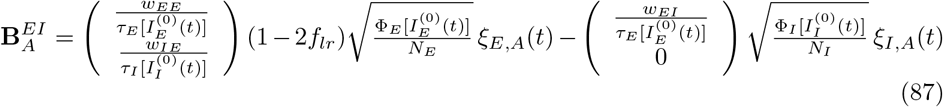

with

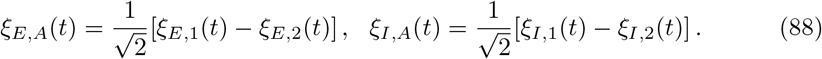

In order to solve Eq. (86), we introduce again the matrix 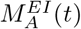, as defined by Eq. (70). The matrix 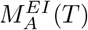 has eigenvectors 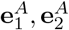 with eigenvalues 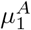 and 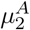. The two eigenvalues and eigenvectors are real when the coupling *f_lr_* between the two modules is sufficiently small (*f_lr_* ≾ 0.13 for our reference parameters, see Fig. 6G), but they can be complex for larger couplings.

Eq. (86) can be explicitely integrated by writing *δ***I**_*A*_(*t*) and **B**_*A*_(*t*) in the basis obtained by translating 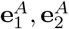 along the limit cycle,

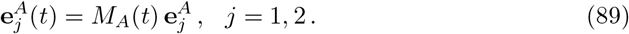

Analogously to a single oscillatory E-I module subject to a forcing current (Eqs. (43,44)), we define

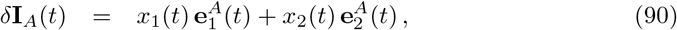

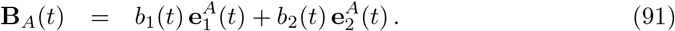

Substitution of these expressions in the differential equation (86) simply gives

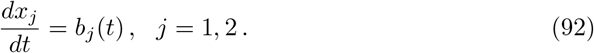

One readily obtains, for 0 ≤ *t* ≤ *T*

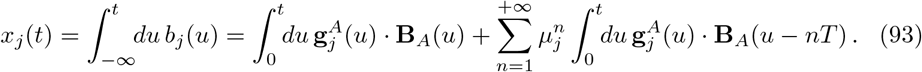

We note that

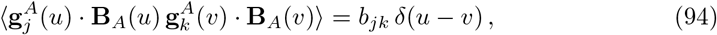

with

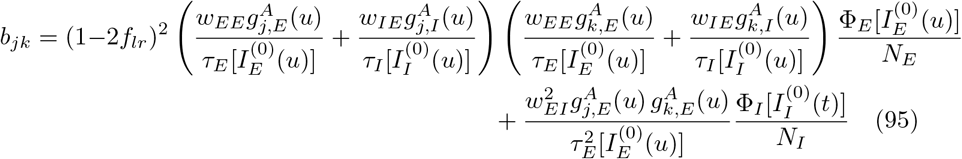

The amplitude of the current fluctuations along the limit cycle is finally obtained, with the help of Eqs. (90), (93), and (94), as

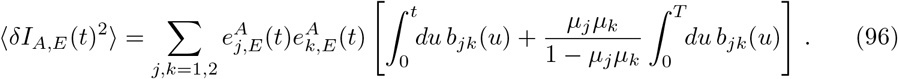

Since 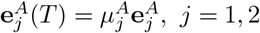, the obtained expression is periodic in time, as it should.

Because we are considering fluctuations around the stationary limit cycle to linear order, the firing rates can be approximated by 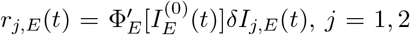, up to the stochastic contribution due to Poisson sampling. The amplitudes of the rate fluctuations 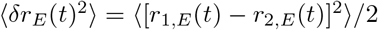 along the limit cycle, up to the direct Poisson contribution, are thus directly obtained by multiplying the above expression with 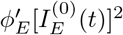. Averaging over one oscillation period, we find

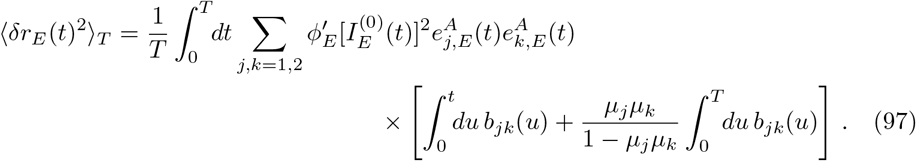

For larger coupling strengths, this result is found to be much more precise when compared to numerical simulations than the expressions obtained in the weak-coupling limit, Eq. (80) or respectively (82) when taking the periodicity of the limit cycle dynamics into account, see Fig. 5. Note however that expression (97) is valid only for small perturbations and fails to account for the limited rate correlations at vanishing coupling due to the periodicity of the limit cycle dynamics, as done in Eq. (82).

In the weak-coupling regime (*f_lr_* → 0), Eq. (97) gives back the simple expression (16) obtained from the stochastic phase equation. This can be seen as follows. When 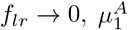 tends towards 1 (Eq. (77)), its value for a single module. Therefore, the r.h.s. of Eq. (97) is dominated by the *j* = *k* =1 contribution in the sum and more precisely by its second term, the denominator of which vanishes. Namely, with the help of Eq. (77), when *f_lr_* → 0 one obtains

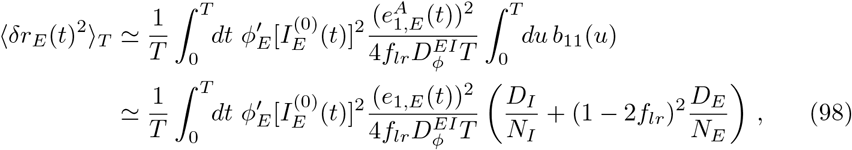

where the last equality is obtained by comparing the definition (95) to the previous expressions for *D_E_* and *D_I_* and noting that the vectors 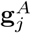 and 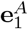 tend toward those of the single module limit cycle, **g**_*j*_ and **e**_1_, when *f_lr_* → 0. Finally, in the limit of weak coupling and weak noise where the phase is well-defined, the mean square difference of excitatory firing rates between the two modules is related to their mean square phase difference,

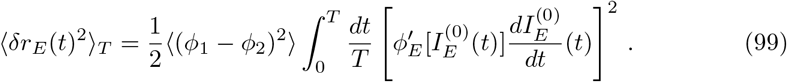

(We used this relation before when deriving Eq. (80).) Comparison of Eq. (98) and Eq. (99) gives back the expression (16) for the mean square phase difference when one remembers that the tangent vector **e**_1_(*t*) is simply the velocity along the current limit cycle.

### Stability analysis of full synchronization for a chain of oscillatory E-I modules

The fully synchronized oscillatory state **I**_*n*_(*t*) = **I**^(0)^(*t*), *n* =1, …, *L*, is also always an exact solution for the dynamics of a chain of *L* E-I modules (Eqs. (17) and (18) or (19) depending on the connectivity). Its stability can be assessed very similarly to the previous case of two coupled modules.

We consider slightly perturbed evolutions for the *L* modules **I**_*n*_(*t*) = **I**^(0)^(*t*) + *δ***I**_*n*_(*t*), *n* = 1, …, *L*. The linear evolution of the perturbations *δ***I**_*n*_ is found to be

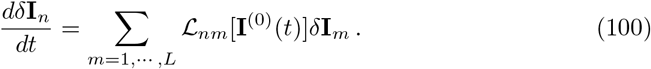

Since the chain is invariant by translation, one can reduce this 2*L*-dimensional evolution to the evolutions of *L* independent 2-dimensional systems. Namely, we write the perturbations as

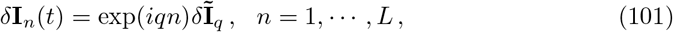

with *L* “wavevectors” *q* = 2*πk/L, n* = 0, …, *L* — 1. This gives, for each *q*, the evolution

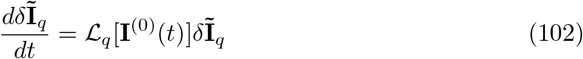

with 2 × 2 matrices 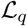 that we give below.

For *E → E* connectivity, the matrices 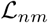 read, for *n = m*,

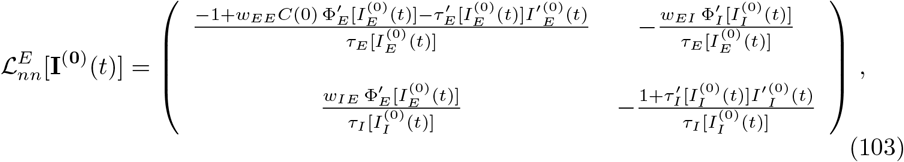

and for *n ≠ m*,

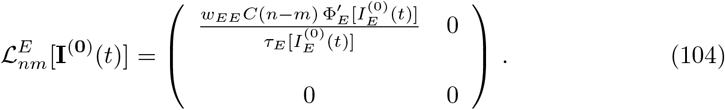

The corresponding matrices 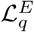 read

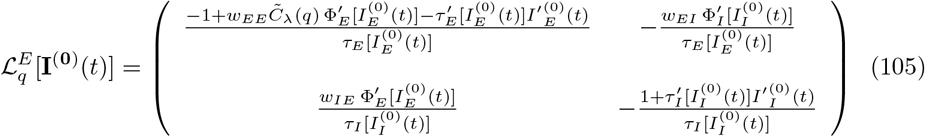

The matrix 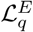 is identical to the previous matrix 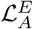 (Eqs. (69,71,72)) governing the stability of two coupled modules with the replacement of (1 — 2*f_lr_*) by the effective coupling 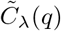,

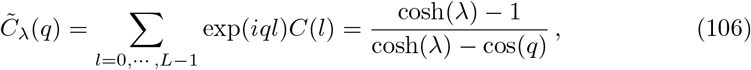

where the second equality specifically holds for the exponentially decreasing excitatory interaction (20). The instability of full synchronization for two-coupled E-I modules for 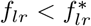 translates into an instability of full synchronization in a chain of modules for close enough to 1. Namely, spontaneous phase gradients appear in the chain of E-I modules for |*q*| < *q*\* with the threshold *q*\* given by Eq. (22) in the main text (for the coupling choice (20)).

For *E → E, I* connectivity, the matrix 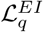 is similarly given by the previous matrix 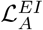 (Eq. (73)) with (1 — 2*f_lr_*) replaced by C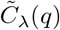. Therefore in this case, the eigenvalues of 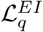 are of magnitude smaller than 1, perturbations of any wavelength tend to vanish, and full synchronization of the chain modules is stable. The evolution of a long wavelength mode can be directly transcribed from our previous results in the two-module case (Eqs. (13) and (15) or Eq. (77)), using the correspondence between *f_lr_* and 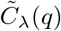,

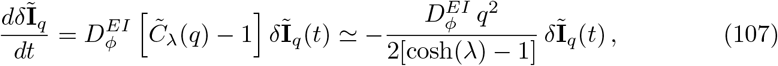

where in the second approximate equality we used that *q* ≪ 1. This provides the relaxation rate of long-wavelength modes given in Eq. (23) of the main text.

### Stochastic dynamics of the phases of oscillations along a chain of E-I modules

We analyze the competition between synchronization and stochastic fluctuation in a chain of E-I modules. We consider *E → E, I* connectivity (Eqs. (17,19)), for which the oscillations of all modules are stably synchronized in the absence of stochastic fluctuations of activity. We first consider the case when the variation of the phase of oscillation is small. The dynamics of these long wavelength modes can be fully described analytically and provide reference expressions. Shorter wavelength fluctuations are considered in the next section.

Taking Eq. (17) as an example, we rewrite the long-range synaptic coupling as

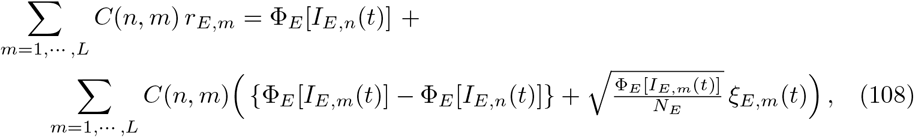

where we have made use of the expression (5) of the firing rate. The term between large parenthesis on the r.h.s. of Eq. (108) can be treated perturbatively when one assumes that the activities of nearby modules are close, and that the noise is not too strong. As before, we suppose that a good starting approximation for the currents **I**_*n*_(*t*) = (*I_E,n_*(*t*), *I_I,n_*(*t*)) which describe the dynamics of the *n*th module is provided by 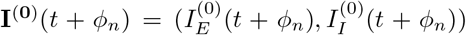, namely the vector of currents on the deterministic limit cycle at a particular phase *ϕ_n_*. Linearization of the chain dynamics around this approximation gives, similarly to Eqs. (38,39),

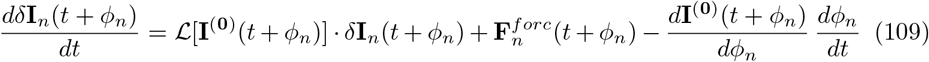

where the matrix 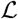 is given by Eq. (41). The forcing of the *n*th module 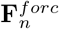 can be decomposed as a sum of inputs coming from the mean activities, **I**^(0)^(*t + ϕ_m_*), of nearby modules and of fluctuations of their activities,

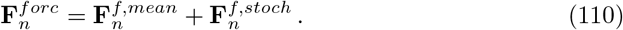

With the help of Eq. (108), one obtains

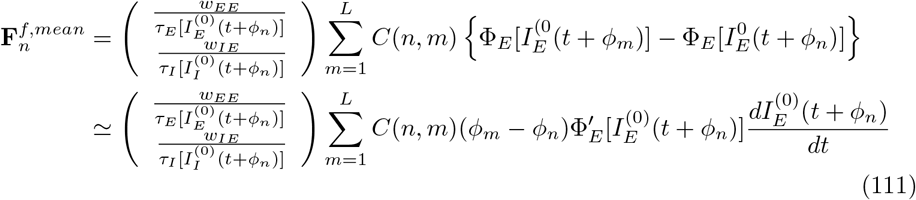

and

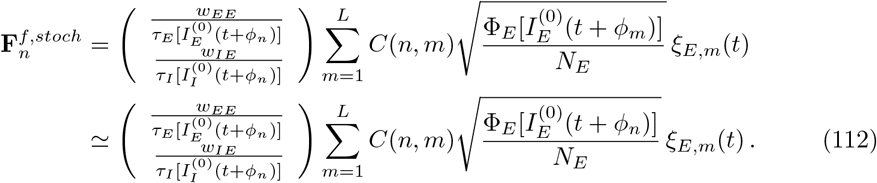

In Eq. (111), we have approximated in a linear way the difference of activities between the modules *n* and *m*. This supposes that the phase difference between the two modules stays close when their distance is small enough for *C*(*n, m*) not to be negligible. (Note that this approximation simplifies the phase equation below but it is not required; without it, the linear phase difference between module n and m would be replaced by a nontrivial function of the phase difference, as in Eq. (60) for the two-module case.) In Eq. (112), we have neglected the phase difference between the two modules in the amplitudes of the stochastic terms since these terms are already treated perturbatively.

Substitution of the forcing term (110) along with the expressions (111,112) in Eq. (50) provides coupled equations for the oscillation phases along the chain,

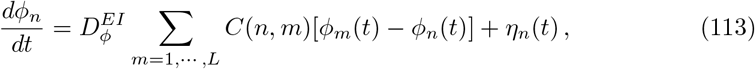

with the constant 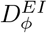 given by Eq. (63). The value of 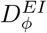 for the reference parameters is provided in Table 1. The stochastic terms *η_n_* in Eq. (113) are Gaussian and are fully characterized by their correlation functions

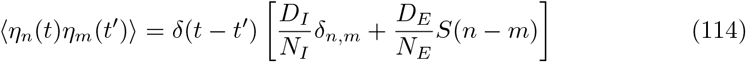

with

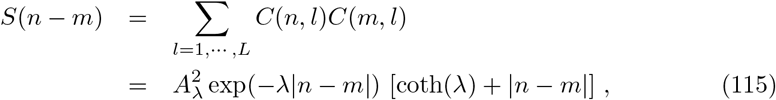

where the second equality in Eq. (115) holds for the exponentially decreasing coupling (20) and where, for simplicity, the expression has been written for a long chain (i.e., neglecting exp(—*L*λ) for *L* ≫ 1/λ). Eqs. (53,54) give the expressions of the constants *D_E_* and *D_I_* that quantify the phase diffusion of an oscillatory E-I module due to stochastic fluctuations.

### Characterization of the E-I module chain stochastic activity profile

#### Long-wavelength modes

As described by Eq. (113), the phases of the different oscillating E-I modules of the spatially extended network are stochastic quantities. Since Eq. (113) is linear and the stochastic terms are Gaussian, the phases also form a Gaussian field that can be fully characterized by its correlation functions. Since the chain is translation invariant, these are most easily computed by writing the phases *ϕ_n_* and the stochastic terms *η_n_* in Fourier space,

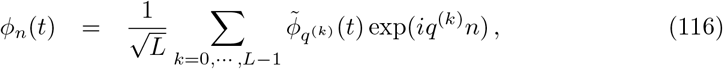

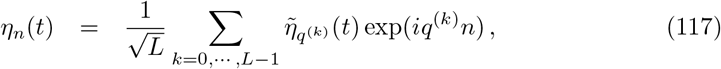

where the *L* wavevectors are {*q*^(*k*)^ = 2*πk/L, k* = 0, …, *L* — 1}. The stochastic terms 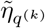 are Gaussian as linear sums of the *η_n_*(*t*) and their correlations are obtained from Eqs. (114, 115) as

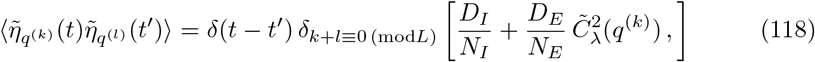

where we have made use of the identity

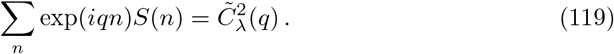

The Fourier transform of the coupling function, 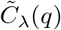, is given in Eq. (106). One can note that its normalization implies that 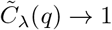 in the long-wavelength limit *q* → 0.

Replacement of expressions (116) and (117) into Eq. (113) gives

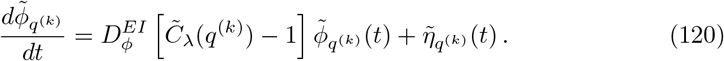

Note that the deterministic part of Eq. (120) exactly corresponds to our previous Eq. (107) for the relaxation of the long-wavelength modes.

For *k* ≠ 0, the above equation can easily be integrated to obtain 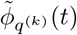,

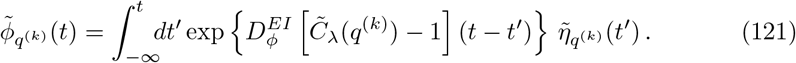

Therefore, the mean square amplitude for the E-I module oscillation phases for a modulation of wavevector *q*^(*k*)^ ≠ 0 is given by

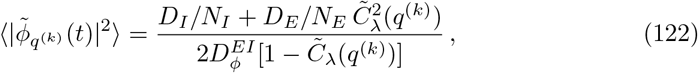

where we have used the expression (118) for the noise correlation. Eq. (122) is the analog for the chain of modules of Eq. (16) in the two-module case. It is relatively straightforward to transform the 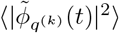 into an expression for the amplitudes of the firing rate fluctuations 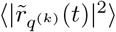 for *k* = 0, along the lines of the previous calculations. Namely, we find

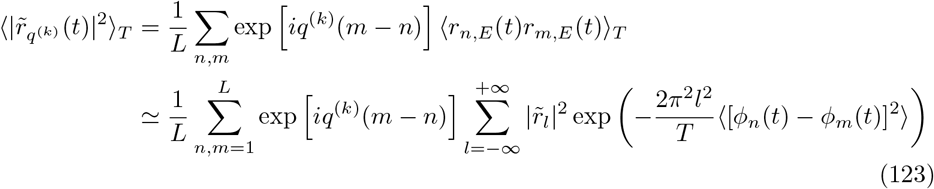

with

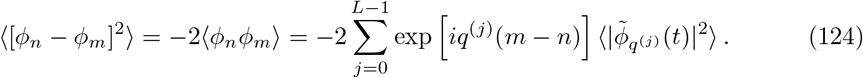

Expression (123) can be expected to hold at large wavelengths, or small *q*, while it will fail at shorter wavelengths where the effective coupling is stronger. It is shown (including an additional contribution 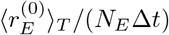 due to Poisson sampling of spikes) alongside a more precise estimate valid for all couplings (see below) and FAT rate model results in Fig. 7I.

When *q* → 0, the coupling strength vanishes. Eq. (122) shows that the mean square amplitudes of the long-wavelength modes then diverge as *q*^-2^,

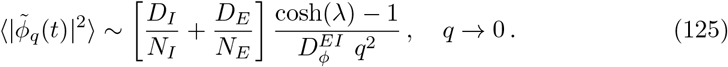

In this limit, the phase equation (121) reduces to the classical Edwards-Wilkinson equation [69],

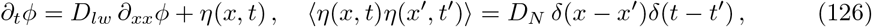

where *D_N_* = *D_I_/N_I_* + *D_E_/N_E_* is the local module noise amplitude (Eq. (9)) and the long-wavelength diffusion constant *D_lw_* is given by

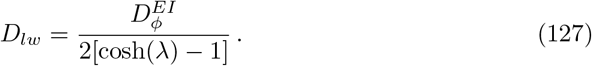

#### Fluctuations of arbitrary wavelengths

Eqs. (113) and (122) very explicitly quantify the dynamics of long-wavelength modes and their average amplitude. Their validity depends however on the fact that for *q* → 0, one eigenvalue of the matrix 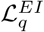 is very close to 1 and the corresponding (phase) mode dominates the fluctuation of activity at the corresponding wavelength 2*π/q*. For modulations of smaller wavelengths, the modules are more strongly coupled. The two modes of 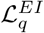 significantly contribute to the fluctuation of activity and Eq. (122) looses its accuracy. In this case, a more precise estimate of the fluctuation amplitude is obtained, for small noise, by linearizing around the fully synchronized state. This extends to a chain of E-I modules our previous analysis for two coupled E-I modules (Eqs. (83)-(96)).

As in the above analysis of the stability of the synchronized state for the chain, we consider perturbed evolutions for the *L* modules around the fully synchronized states **I**_*n*_(*t*) = **I**^(0)^(*t*) + *δ***I**_*n*_(*t*), *n* =1, …, *L*. The linear evolution of the perturbations *δ***I**_*n*_ is found to be

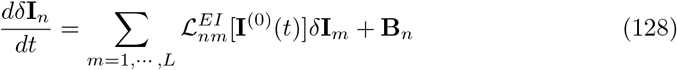

with the stochastic terms **B**_*n*_ given by,

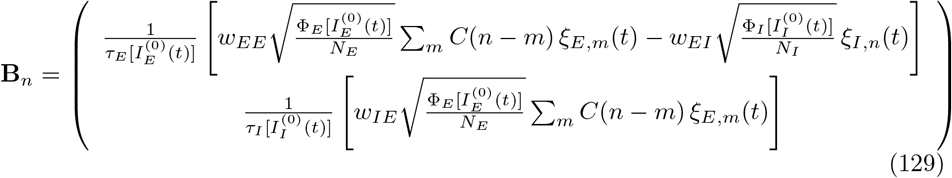

As above (Eq. (101),(117)), translation invariance allows us to simplify the coupled equations (128) by introducing the Fourier representations

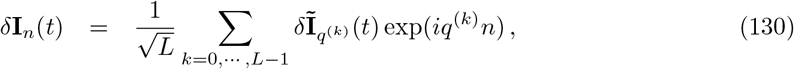

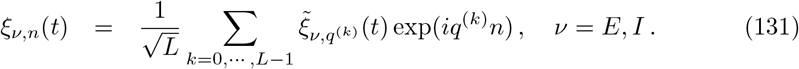

The noise Fourier components obey,

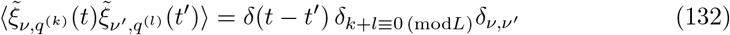

Replacement of Eqs. (130,131) in Eq. (129) gives decoupled dynamics for each Fourier mode of the currents 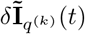, generalizing Eq. (102) to account for stochasticity,

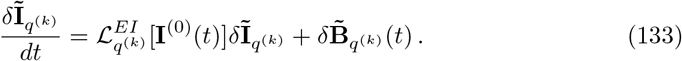

The stochastic terms 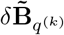 are the Fourier components of the **B**_*n*_ (Eq. (129)),

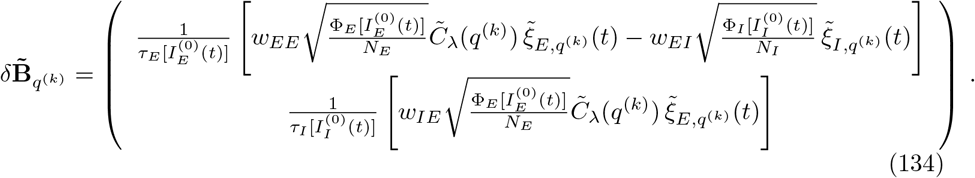

Eq. (133) and (134) generalize to arbitrary Fourier modes the previous Eq.(86) and (87) for the antisymmetric mode in the two-module case with the with the already-noted replacement of (1 — 2*f_lr_*) by the Fourier transform 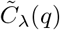 (Eq. (106)). The amplitudes of the Fourier mode modulations thus read, similarly to Eqs. (95,96),

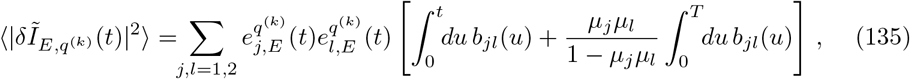

where the vectors 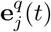 are defined for the matrices 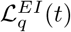 as the vectors 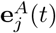 for the matrix 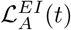.

The amplitudes of the rate fluctuations are again obtained by multiplying the expression above by 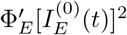, and averaged over the complete limit cycle we find

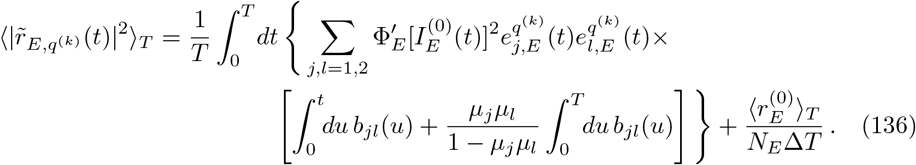

Here, we explicitly added the contribution that arises due to Poissonian sampling of the spiking dynamics within bins of size Δ*T*. The above result is compared to the stochastic FAT rate model in Fig. 7I.

## Supporting information

**S1 File. Tabulated values for** Φ_*σ*_ (*I*) **and** *τ*_(*fat*)_(*I*s).

**S1 Fig. The FAT rate model for EIF neurons.** Plots of the f-I curve and the fitted adaptive timescale (FAT) used throughout the manuscript, and an assessment of the FAT rate model’s capacity to account for time-varying input current.

**S2 Fig. Oscillatory E-I module dynamics for reference parameters B and C.** Comparison of spiking network and FAT rate model dynamics for different synaptic strengths that those used throughout the manuscript.

**S3 Fig. Dynamical regimes of two coupled E-I modules for reference parameters B and C.** Phase diagrams of the dynamical regimes of two E-I modules coupled by long-range *E → E* excitatory connectivity, for different synaptic strengths that those used throughout the manuscript.

**S4 Fig. Average phase gradients for a chain of E-I modules.** The probability distribution of extended phase gradients 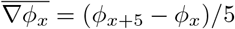 that exceed the standard deviation of locally contributing phase differences 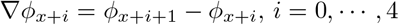.

**S5 Fig. Phase gradients in a chain of E-I modules with decreased long-range excitation.** Histograms of local phase differences and extended phase gradients for different space decay constants λ and network sizes *N* (as well as both types of long-range connectivity studied), when the contribution of long-range excitation to the local dynamics is diminished by a factor 2.

## Acknowledgments

We would like to thank Joran Deschamps for performing preliminary rate-model simulations and computations on synchronization of E-I modules during an M2 internship. We are also grateful to Frédéric Chavane and Thomas Brochier for instructive comments as well as to Josef Ladenbauer and Nicolas Brunel for drawing our attention respectively to ref. [50] and ref. [13]. We also wish to thank Bard Ermentrout for drawing our attention to ref. [22] which had escaped our notice. AK acknowledges the support of Fondation de la Recherche Médicale (FRM) through a doctoral FRM fellowship.

---

**Fig S1.**
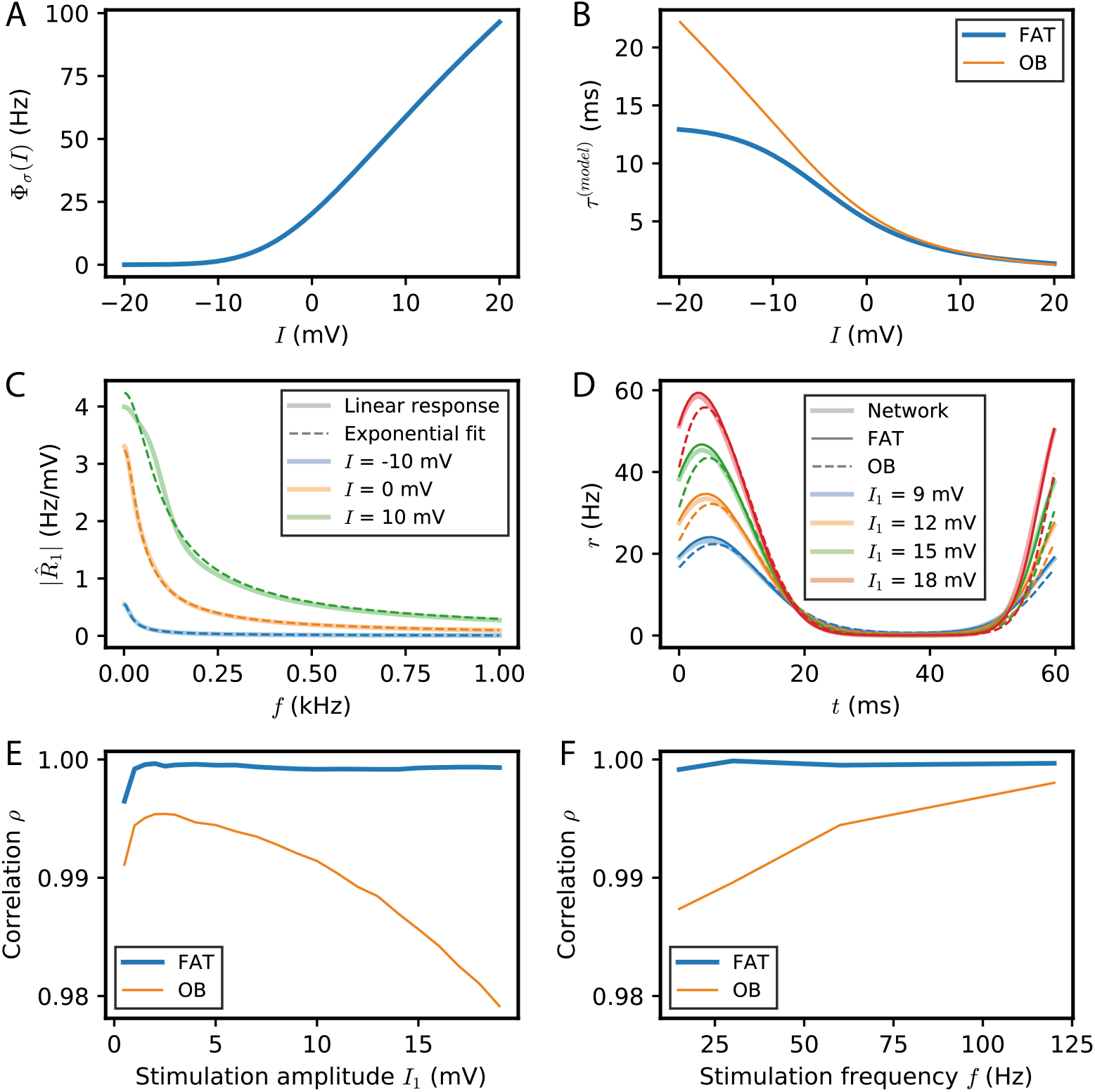
The FAT rate model for EIF neurons. (A) The f-I curve Φ_*σ*_(*I*) for the EIF neurons used in the rate-model formulation (numerically evaluated following [71]). (B) The fitted adaptive timescale *τ*^(*FAT*)^(*I*) determined by fitting the linear firing-rate response and used throughout the manuscript (thick blue line). For comparison, we also show the adaptive timescale proposed by Ostojic and Brunel (OB) [48] (thin orange line). (C) Examples of the fit of the analytically determined firing-rate response with the Fourier transformation of an exponential kernel in the time domain (see *Methods*). (D) Dynamics of a population of (uncoupled) EIF neurons and the two adaptive rate models upon injection of a sinusoidal current *I*_1_ sin(2*πft*) for *f* = 17 Hz and different values of *I*_1_. (E,F) Correlation between the network activity and the two adaptive rate models (FAT, thick blue; OB, thin orange) as a function of the amplitude *I*_1_ of the injected current for a fixed frequency *f* = 17 Hz (E), and as a function of frequency *f* for a fixed amplitude *I*_1_ = 3 mV (F).

**Fig S2.**
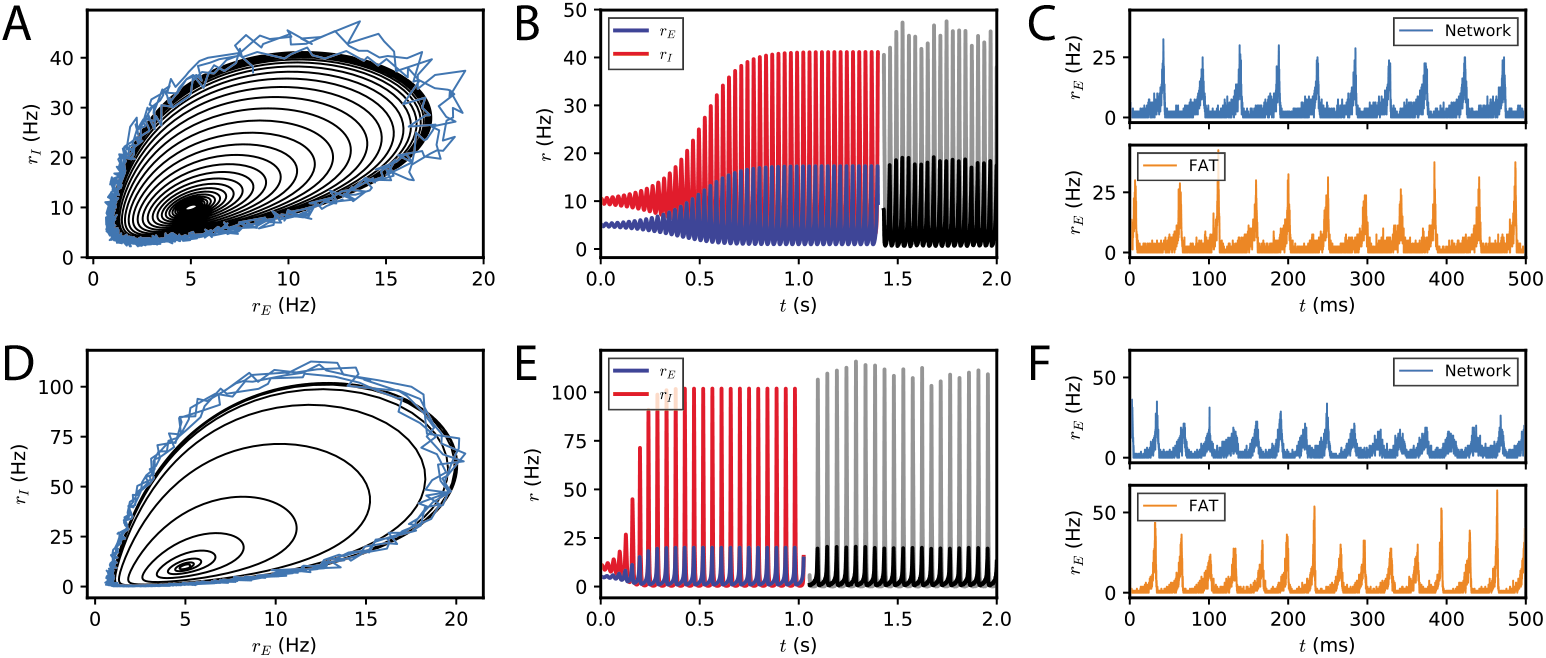
Oscillatory E-I module dynamics for reference parameters B and C. (A,B) The deterministic limit cycle dynamics for a single E-I module described by the FAT rate model with synaptic strengths corresponding to reference case B (see Fig. 1A) is shown together with spiking network simulations with *N* = 10^6^. The synaptic strengths are given by *w_EE_* = 1.6mVs, *w_IE_* = 1.6 mVs, and *w_EI_* = 0.8 mVs. (C) A comparison of the stochastic FAT rate model with spiking network simulations with *N* = 10^4^ for the same parameters. (D-E) Analogous plots for the reference case C, with *w_EE_* = 1.76 mVs, *w_IE_* = 3.2 mVs, and *w_EI_* = 0.4 mVs.

**Fig S3.**
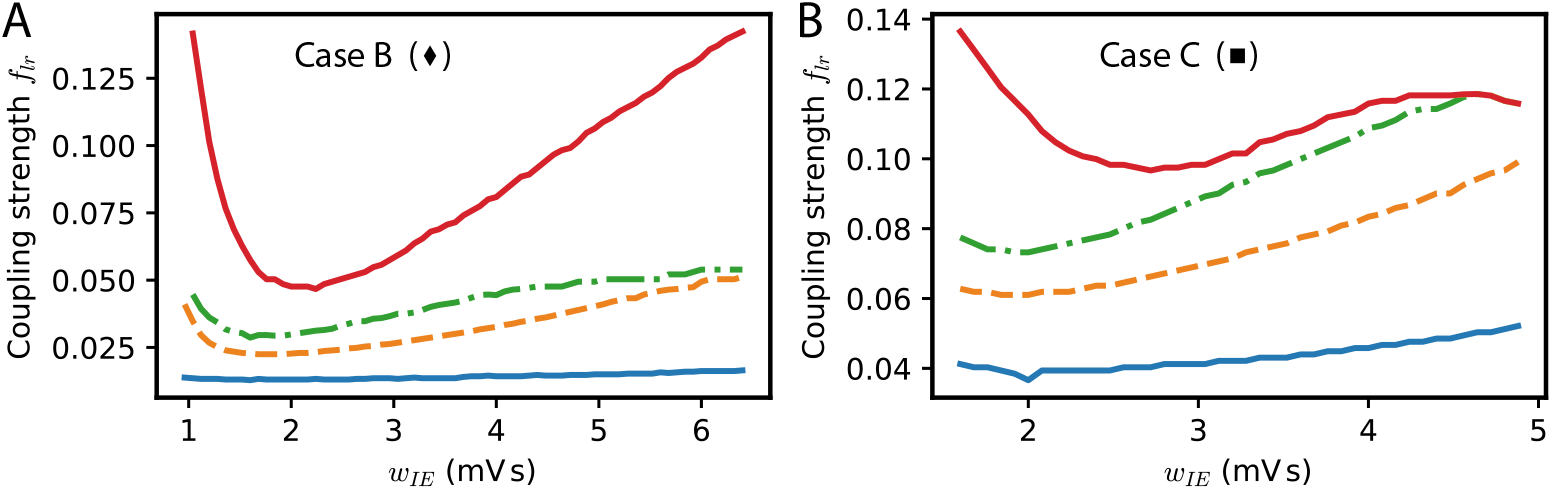
Dynamical regimes of two coupled E-I modules for reference parameters B and C. The analog of Fig. 2B for the two other reference parameters shown on Fig. 1A. The phase diagrams for case B (*w_EE_* = 1.6 mVs, *w_IE_w_EI_* = 1.28 mV^2^s^2^) and case C (*w_EE_* = 1.76 mVs, *w_IE_w_EI_* = 1.28 mV^2^s^2^) are shown on panels (A) and (B), respectively.

**Fig S4.**
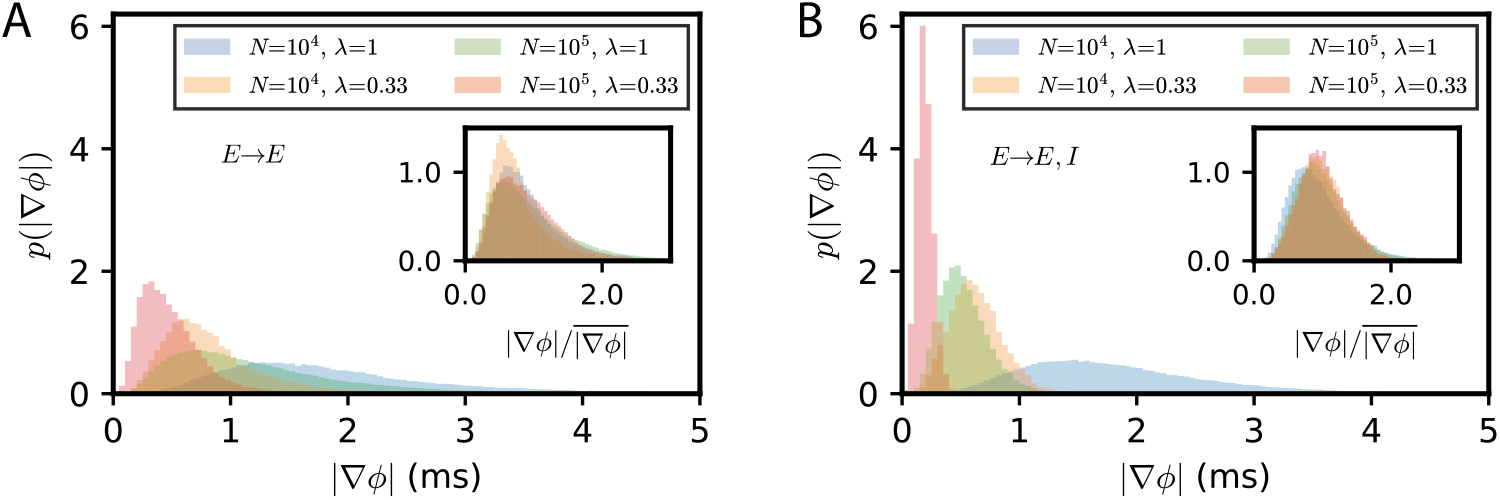
Average phase gradients for a chain of E-I modules. The probability distribution of extended phase gradients 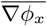 that exceed the standard deviation of locally contributing phase differences (see *Methods*), for different values of λ and *N*, for *E → E* connectivity (A) and *E → E,I* connectivity (B).

**Fig S5.**
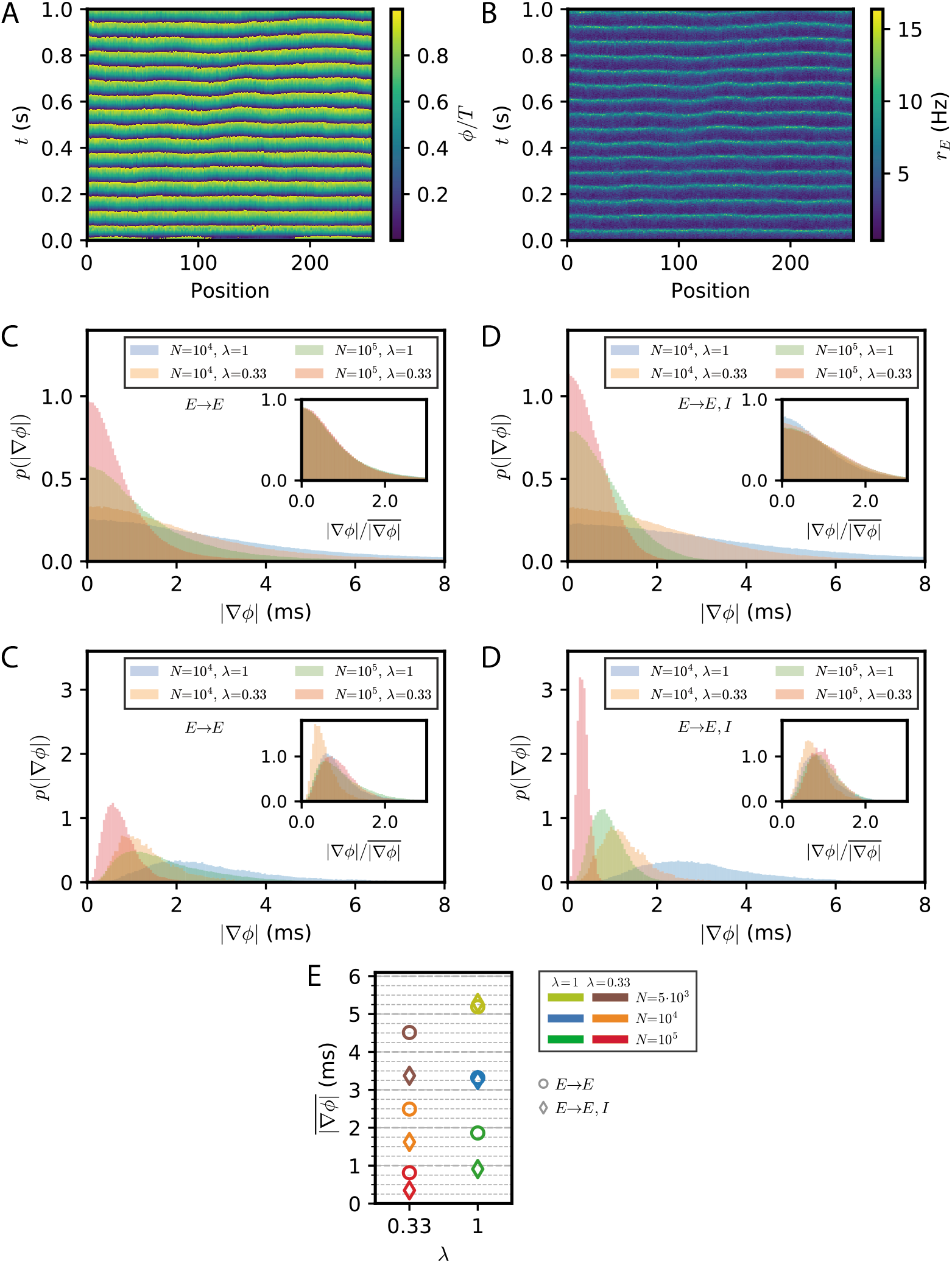
Phase gradients in a chain of E-I modules with decreased long-range excitation. (A,B) Simulation of the stochastic rate model (FAT) for a chain with *E → E, I* connectivity, a space constant λ = 0.33, and network size *N* = 10^4^. In contrast to Fig. 7D,E, the contribution of the long-range connectivity to the total recurrent excitatory drive is lowered by a factor 2 at the expense of purely local recurrent excitation. (A) Instantaneous phase as obtained from the Hilbert transform of bandpass-filtered excitatory activity, where the (unfiltered) rates are shown in (B). (C,D) Histograms of the local phase differences for different values of λ and *N* for *E → E* connectivity (C) and *E → E, I* connectivity (D). (E,F) Histograms of the extended phase gradients 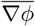. The averages of the distributions shown in (E,F) are plotted in (G), and are the analog of Fig. 7I for a diminished contribution (by a factor of 2) of long-range connectivity to the total recurrent excitatory drive.

